# Danicamtiv increases myosin recruitment and alters the chemomechanical cross bridge cycle in cardiac muscle

**DOI:** 10.1101/2023.01.31.526380

**Authors:** Kristina B. Kooiker, Saffie Mohran, Kyrah L. Turner, Weikang Ma, Galina Flint, Lin Qi, Chengqian Gao, Yahan Zheng, Timothy S McMillen, Christian Mandrycky, Amy Martinson, Max Mahoney-Schaefer, Jeremy C. Freeman, Elijah Gabriela Costales Arenas, An-Yu Tu, Thomas C. Irving, Michael A. Geeves, Bertrand C.W. Tanner, Michael Regnier, Jennifer Davis, Farid Moussavi-Harami

## Abstract

Modulating myosin function is a novel therapeutic approach in patients with cardiomyopathy. Detailed mechanism of action of these agents can help predict potential unwanted affects and identify patient populations that can benefit most from them. Danicamtiv is a novel myosin activator with promising preclinical data that is currently in clinical trials. While it is known danicamtiv increases force and cardiomyocyte contractility without affecting calcium levels, detailed mechanistic studies regarding its mode of action are lacking. Using porcine cardiac tissue and myofibrils we demonstrate that Danicamtiv increases force and calcium sensitivity via increasing the number of myosin in the “on” state and slowing cross bridge turnover. Our detailed analysis shows that inhibition of ADP release results in decreased cross bridge turnover with cross bridges staying on longer and prolonging myofibril relaxation. Using a mouse model of genetic dilated cardiomyopathy, we demonstrated that Danicamtiv corrected calcium sensitivity in demembranated and abnormal twitch magnitude and kinetics in intact cardiac tissue.

**Significance Statement:** Directly augmenting sarcomere function has potential to overcome limitations of currently used inotropic agents to improve cardiac contractility. Myosin modulation is a novel mechanism for increased contraction in cardiomyopathies. Danicamtiv is a myosin activator that is currently under investigation for use in cardiomyopathy patients. Our study is the first detailed mechanism of how Danicamtiv increases force and alters kinetics of cardiac activation and relaxation. This new understanding of the mechanism of action of Danicamtiv can be used to help identify patients that could benefit most from this treatment.

## Introduction

The prevalence of heart failure (HF) continues to rise and despite new and improved treatments, there is still significant morbidity and cost (1). More than half of the HF patients can be categorized by depressed systolic function or contractility (2). Traditional inotropic agents improve cardiomyocyte contractility via increased intracellular cyclic adenosine monophosphate (cAMP) and calcium levels but do not improve patient survival (3, 4). Directly targeting sarcomeric proteins can overcome adverse effects of traditional inotropes such as arrythmias, increased myocardial oxygen demand and activation of cell death pathways (3). Small molecules targeting myosin and modulating its activity have been developed over the last decade for treatment of HF. The strategy for identifying these compounds has been to identify molecules that selectively bind to cardiac myosin and increase or inhibit the myosin ATPase activity. Omecamtiv Mecarbil (OM) is a first in-class myosin activator that was originally thought to work via increased phosphate release and priming myosin for binding to actin (5, 6). More detailed biophysical measurements showed that OM actually inhibits the myosin working stroke and prolongs the detachment of a small population of non-force generating myosin heads (7, 8). The OM-bound myosin heads activate the thin filament and recruit more myosin heads with the net effect of having more myosin heads that are available for force generation. This mechanism relies on myosin co-operativity and is dose dependent with force inhibition at higher doses both *in vitro* and *in vivo.* OM recently completed phase III clinical trial in heart failure and was found to result in a modest decrease in heart-failure events or death from cardiovascular causes compared to placebo in particular with patients with lowest ejection fraction deriving the most benefit (9, 10). The successes and challenges of OM show the importance of better understanding of the molecular mechanisms of new sarcomere modulators.

Danicamtiv (Dani) or MYK-491 is a promising novel myosin activator that increases myofibril ATPase activity and calcium sensitivity (11). *In vivo* studies demonstrated improved atrial and ventricular function in a canine HF model as well as patients with heart failure with reduced ejection fraction that were enrolled in a phase 2a randomized clinical trial (11). There is an ongoing clinical trial enrolling patients with genetic dilated cardiomyopathy (DCM) with sarcomeric variants for treatment with Dani. We and others have shown that reduction in cardiac twitch, the tension developed as a function of time is predictive of diagnosis and severity of DCM in rodent myocytes and humans (12–14). It is logical that DCM patients may benefit from treatment with myosin activators. A recent study in a human engineered heart tissue platform showed that administration of Dani resulted in a larger increase in systolic contraction at a smaller lusitropic cost compared to OM (15). While this suggests possible differences in mechanisms, there is limited information available regarding Dani’s mechanisms of action other than that Dani increases the number of myosin heads available for force production without altering passive stiffness (16). Here, we present the first detailed mechanistic study of how Dani affects the cross bridge cycle, cardiac muscle contractile kinetics and myosin structure. We show that Dani increased calcium sensitivity and elevated force mostly at low calcium levels. Using X-ray diffraction, we demonstrate that treatment with Dani under resting conditions repositions myosin heads closer to the thin filament. In myofibrils under load, Dani slowed the myosin cross bridge cycle and prolonged relaxation kinetics by inhibiting the rate of ADP product release. Lastly, we demonstrate the ability of Dani to recover normal calcium sensitivity and twitch force in a thin filament rodent DCM model.

## Results

### Danicamtiv increased submaximal force and calcium sensitivity while prolonging myofibril relaxation

Permeabilized porcine ventricular muscle preparations showed a left-ward shift in the force vs. calcium (pCa) relationship in the presence of 1 μm Dani (Figure 1A). This was quantified by an increase in pCa_50_ (Figure 1B; 5.72±0.02 vs. 5.91±0.05, p<0.05) indicating an increase in the sensitivity of the myofilament to calcium. While there was a slight increase in maximally activated force (Figure 1C), there was a larger and more significant increase in force at submaximal calcium levels (Figure 1A). This leads to a reduction in the Hill coefficient (Figure 1D; 3.96±0.31 vs. 2.27±0.29, p<0.001), suggesting an effect of Dani on myofilament cooperativity or recruitment. Force redevelopment rate constants(*k*_tr_) were significantly slower at each level of force (Supplemental Figure 1B) suggesting that Dani reduced cross bridge cycling kinetics. Incubation in Dani resulted in more than a 50% decrease in maximal *k*_tr_ (Figure 1E; 1.91±0.14 vs. 0.83±0.08, p<0.01). High-frequency small amplitude oscillation measurements of steady state stiffness showed that Dani increased stiffness to the greatest degree at lower calcium levels (Figure S1C). The increased stiffness is proportional to the increase in force, suggesting the increased stiffness is due to more myosin heads rather than an increase in force per myosin head.

**Figure 1.**
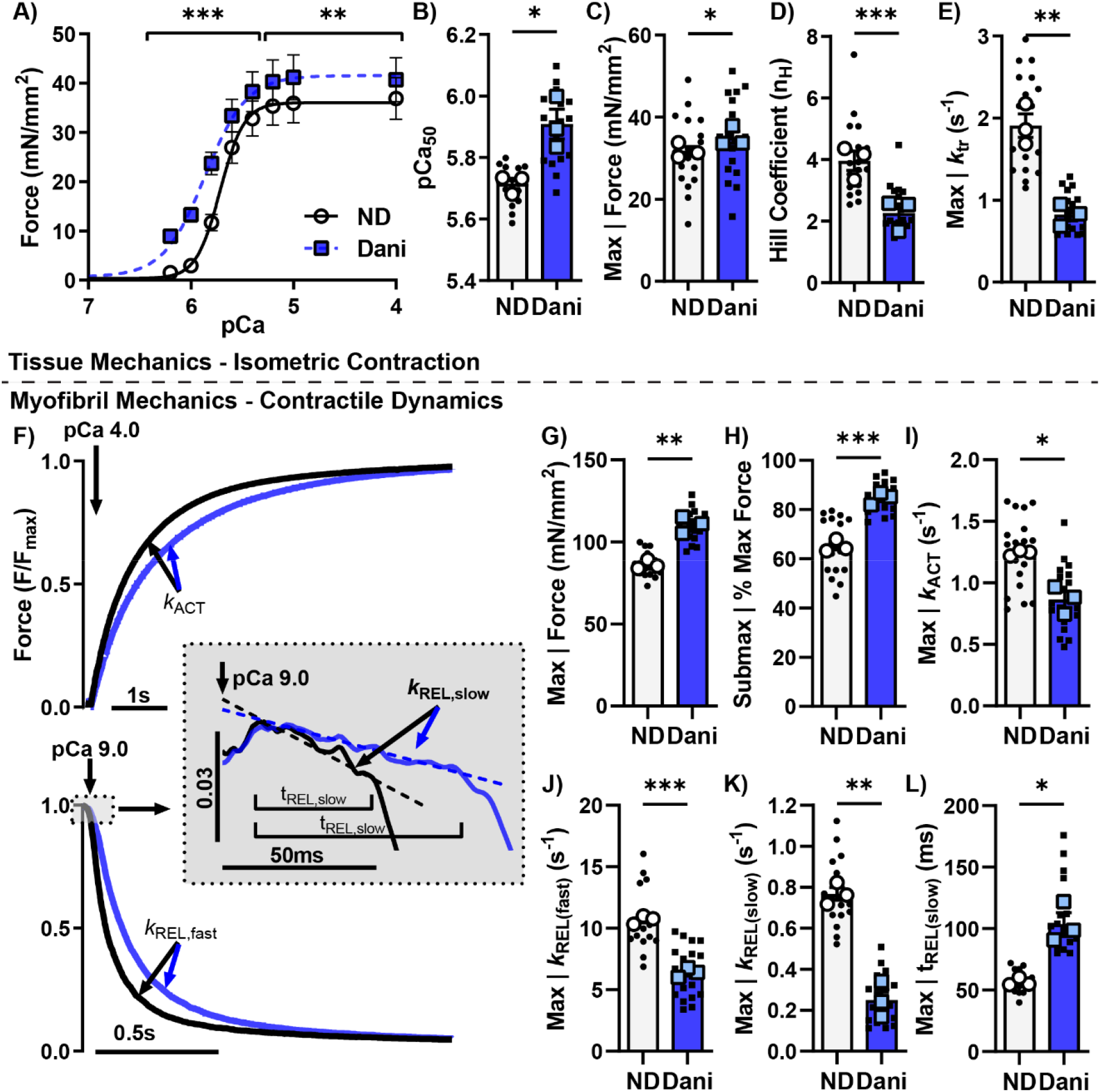
Mechanics in permeabilized β-myosin porcine cardiac tissue and myofibrils treated with 1 μM Danicamtiv demonstrating increased force and calcium sensitivity with prolonged relaxation. Isolated uniaxially aligned muscle strips from porcine ventricle tissue showed a left shift and increased force vs ND in the F-pCa curves (A) after incubation in 1 μM Dani. There was a significant increase in pCa_50_ (B) and max force generation (C) with a significant decrease in Hill coefficient (D) and max *k*_TR_ (E). In myofibril mechanics (F) at maximum calcium (pCa=4.0), we again saw increased force generation (G), with increased submax force production (H). Exponential activation (I) and relaxation kinetics were reduced (J), along with significant decrease in the rate (K) and duration (L) of the slow phase of relaxation. Data is from 19 paired demembranated tissue preparations and 18 DMSO or 21 Dani treated myofibrils from 3 biological replicates. *P < 0.05., **P<0.005, ***P<0.001 vs ND, error bars represent SEM.

Isolation of sub-cellular myofibrils allow for measurements of force and kinetics of both activation and relaxation (17, 18). Figure 1F shows representative normalized force traces of porcine myofibrils transitioning between maximally activated (high Ca^2+^) and relaxed (low Ca^2+^) conditions by rapid solution switching via a double-barreled pipette with and without Dani. Force is developed exponentially with a rate constant (*k*_ACT_) that depends on thin filament activation, myosin recruitment, and myosin cross bridge cycling. The rapid switch back to low calcium solution results in a biphasic relaxation relationship with an initial linear phase followed by rapid exponential phase back to baseline. The rate constant during the linear slow phase of relaxation (*k*_REL(slow)_) reflects the rate of cross bridge detachment, which is independent of the Ca^2+^ dissociation from troponin C. The duration of the slow phase (t_REL(slow)_) reflects the time it takes for the thin filament to deactivate and has been shown to be influenced by the properties of the thin filament proteins and the level of calcium (19, 20). The subsequent fast exponential phase of relaxation (*k*_REL(fast)_) reflects inter-sarcomere dynamics and involves both active and passive elements that are influenced by multiple factors.

As in the demembranated tissue preparations, Dani increased myofibril force at maximal (pCa=4) and submaximal calcium (pCa=5.8). The ratio of submaximal to maximal activation was also significantly increased with Dani. (Figure 1G and H). Treatment with Dani resulted in a slight decrease in *k_ACT_* (Figure 1I). Relaxation kinetics analysis (Figure 1J) revealed that Dani decreased the fast rate of myofibril relaxation (*k*_REL(fast)_; 10.64±0.20 vs. 6.34±0.22 s^-1^, P<0.001), which is consistent with the slowed cellular and organ level kinetics in pre-clinical models (21). Our analysis also revealed that the initial linear slope (*k*_REL(slow)_; 0.77±0.03 vs. 0.25±0.05 s^-1^, P<0.005) of myofibril relaxation was decreased while the thin filament deactivation duration (t_REL(slow)_; 56.5±1.9 vs. 103.7±9.4 ms, P<0.05) increased (Figure 1K and L). These results showing prolonged relaxation kinetics suggest that Dani impacts cross bridge turnover kinetics by inhibiting product release, either phosphate (P_i_) or ADP. Previously reported mechanisms of the myosin activator OM showed that slower cross bridge turnover can result in increased force generation and prolonged myosin head activation, requiring more time for the thin filament to transition from an active to an inactive state (7, 22, 23). The rate of cross bridge turnover has been previously identified as a critical regulator for transitioning the thin filament from an active to an inactive state (24, 25).

While it is known that Dani increases myofibril ATPase rate (11), a detailed understanding of the exact mechanisms is lacking. Together, our tissue mechanics and myofibril measurements suggest that Dani impacts cardiac function through increased myosin head recruitment and altered cross bridge cycling kinetics (Figure 3). We will investigate this further through the following reductionist assays.

**Figure 2.**
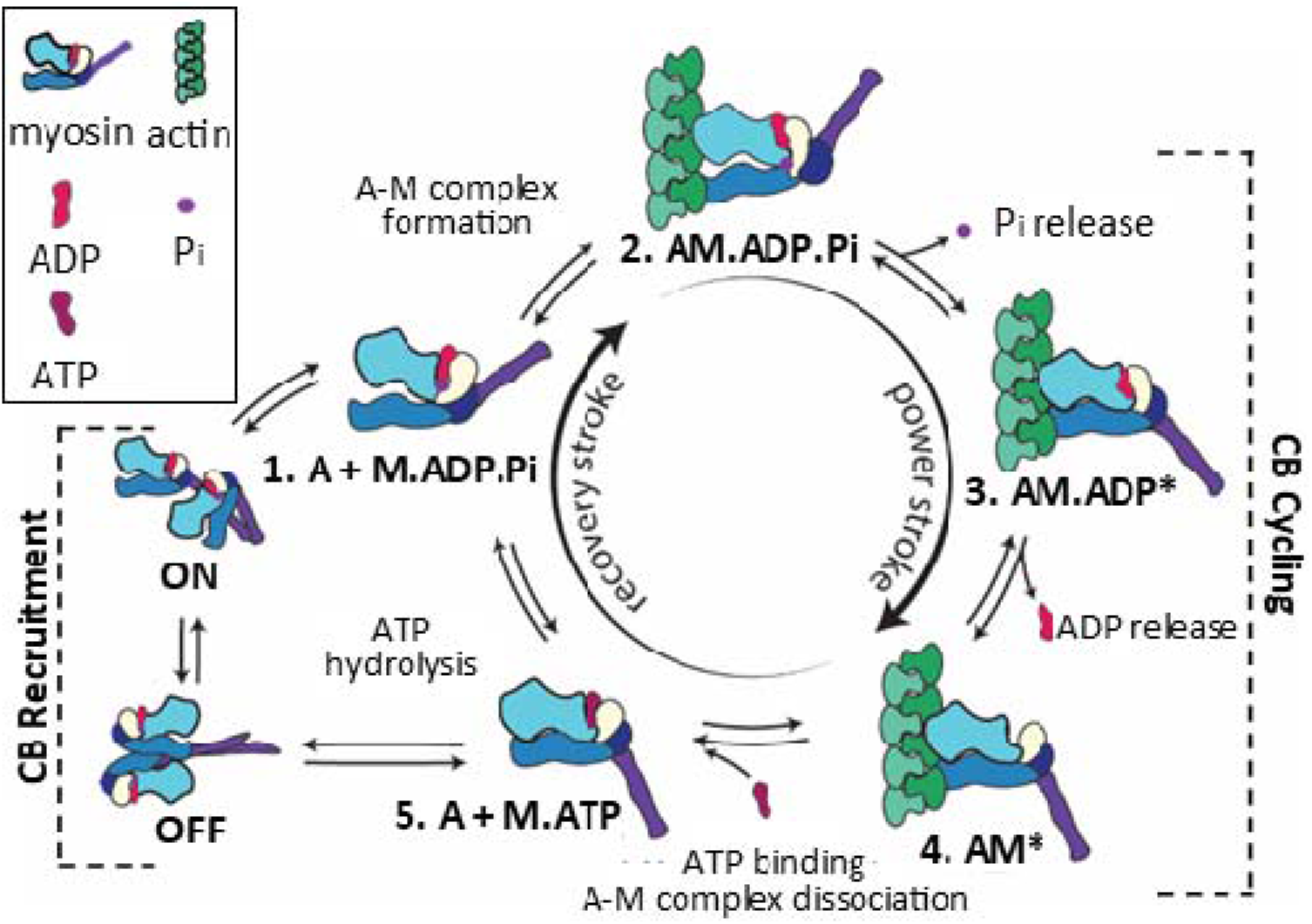
Scheme illustrating a five-step cross-bridge model of contraction. Figure art by Matthew Childers, Regnier Lab, University of Washington, 2023. This work is licensed under a Creative Commons Attribution-NonCommercial 4.0 International License.

**Figure 3.**
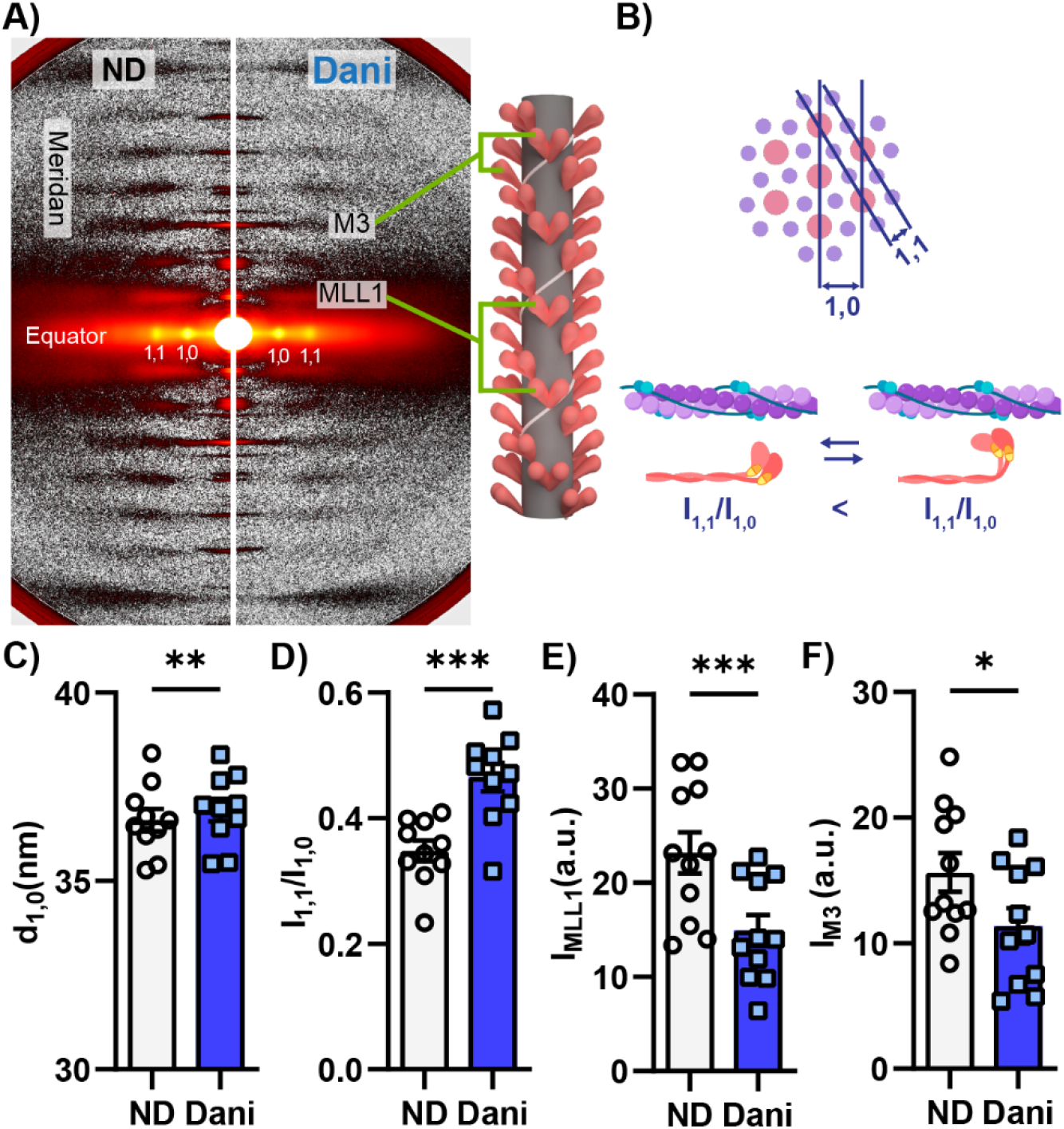
50 μM Danicamtiv partially activated the thick filament in resting cardiac muscle (pCa=8.0). A) Sample X-ray diffraction patterns of demembranated porcine cardiac tissue exposed to X-ray beams while bathed in low calcium solution (pCa=8.0) containing DMSO (left) or Dani (right). In each diffraction pattern, the reflections of interest are labeled and. B) Schematics defining how actin and myosin relationships relate to equatorial measurements. Dani increased the lattice spacing (C, d_1,0_), intensity ratio of the primary equatorial reflections (D, I_1,1_/I_1,0_), intensity of the first-order myosin-based layer line (E, I_MLL1_), and the third-order myosin-based meridional reflection (F, I_M3_). Data is from 10 paired tissue preparations. *P < 0.05., **P<0.005, ***P<0.001 vs. ND, error bars represent SEM, a.u., arbitrary units. The illustrations of lattice spacing diagram and intensity ratio visuals were created with BioRender.com

### Danicamtiv altered resting myosin thick filament structure and both passive and active elastic and viscous moduli

Previous studies have shown that myosin modulators can affect myosin structure (26, 27). We used small-angle-x-ray diffraction patterns from permeabilized porcine myocardium to study the structural changes induced by Dani under relaxed conditions (pCa8). Figure 2A shows representative two-dimensional X-ray diffraction patterns from relaxed permeabilized porcine cardiac preparations in the absence (left panel) or the presence of Dani (right panel). The lattice spacing (d_1,0_; Fig 2B, top), directly proportional to interfilament spacing (28), increased after Dani treatment (Fig 2C). The increase of d1,0 could be a result of increased electrostatic repulsion between the myofilaments when myosin heads move away from the thick filament backbone towards actin filaments (29). The equatorial intensity ratio (I_1,1_/I_1,0_) reflects the proximity of myosin heads to the thin filament (Figure 2B, bottom) (26–28). As seen in Figure 2D, Dani significantly increased I_1,1_/I_1,0_ under relaxed conditions suggesting that myosin heads moved away from the thick filament backbone and closer to the thin filament.

The meridional reflections arise from axially repeating structures in the myofilaments (Figure 2A) (28). The distance of these reflections to the beam center are inversely related to the spacing of the axial periodicities along the myofilaments. When cardiac muscle was exposed to Dani under relaxing conditions, the intensity of the first-order myosin-based layer line (I_MLL1_; Figure 2E) and the third-order myosin-based meridional reflection (I_M3_; Figure 2F), both of which correlate with the ordering of myosin heads on the thick filament backbone (30, 31), decreased by 27% and 35%, respectively. The decrease of I_M3_ and I_MLL1_ reflects a reduction of the number of ordered myosin heads. The intensity of the sixth-order myosin-based meridional reflection (M6) arises primarily from structures within the thick filament backbone (28). The spacing of the M6 reflection (S_M6_) increased by 0.2% with Dani (Figure S2). Both the loss of the helical ordering of the myosin heads and the increase in thick filament backbone periodicity are suggestive of a transition in myosin structure and its position on the thick filament from the OFF to ON state (32). It is well recognized that resting thick filaments can be characterized by different structural and biochemical states that can be affected by both disease and small molecules (27, 33, 34). Here, the increase in number of ON state thick filaments in the presence of Dani contributes to increased force generation via more myosin motors available for contraction (26).

Our x-ray diffraction findings were supported by performing sinusoidal length-perturbation analysis to porcine cardiac tissue with and without Dani under resting and activating conditions. The resultant elastic and viscous moduli responses provide insight into the number of bound cross bridges and characteristics such as kinetics of force generating cross bridges, respectively. As seen in Figure S3, under relaxed conditions (pCa 8.0, 5 mM [MgATP]), elastic moduli were not significantly changed by Dani. There was a significant increase in the viscous moduli at a sub-set of oscillatory frequencies for the Dani-treated strips. Under maximally activated conditions (pCa 4.8, 5 mM [MgATP]) there was an increase in visco-elastic myocardial stiffness for the Dani-treated strips vs. the untreated. Both elastic and viscous moduli values were greater in the presence of Dani across a wide range of frequencies, suggesting greater cross bridge binding in the presence of Dani. The observed leftward-shift towards lower frequencies for the elastic and viscous moduli responses also suggests Dani slowed cross bridge cycling, consistent with the slowed *k*_tr_ reported above (Figure 1E).

### Prolonged myofibril relaxation induced by Danicamtiv was due to altered ADP release rate

Using a combination of reductionistic approaches, we investigated whether Dani affects the product release rates of inorganic phosphate (Pi) and ADP during cross bridge cycling. Either of these mechanisms could explain the decreased crossbridge turnover time suggested by the decreased myofibril relaxation kinetics.

It is well accepted that increasing Pi results in decreasing force (35, 36). As seen in Figure S4, a linear dependence of relative force on Log [P_i_] was found in demembranated porcine cardiac tissue. The difference between the slopes was not significant between the Dani and no drug group. Hence, the relation between force and [P_i_] was unchanged with Dani. These results are different than what is reported for OM, where 1μM OM counteracts the inhibitory effects of P_i_ under similar conditions (37).

To test the hypothesis that ADP release is affected by Dani, we quantified myosin cross bridge activity in the presence of elevated ADP conditions. Initial experiments utilized the *in vitro* motility (IVM) assay, which directly measures cross bridge activity, and assessed how elevated Dani impacted purified actin and myosin interactions compared to untreated and elevated ADP conditions (Figure 5A). Previous studies have shown that when ADP levels are elevated, filament velocity is decreased because ADP release is inhibited, promoting the strongly bound actin-myosin state (38, 39). As seen in Figure 5B, Dani also resulted in a substantial (~55%) decrease in filament velocity (2.65±0.17 vs. 1.13± 0.12 μm/s). There was no further decrease in velocity with nucleotide mixtures containing 50% ATP and 50% ADP in the presence of Dani (1.01± 0.05 μm/s, p=0.96), suggesting no additive effect. These results support the hypothesis that Dani slows cross bridge cycling rate through slowed ADP release.

**Figure 4.**
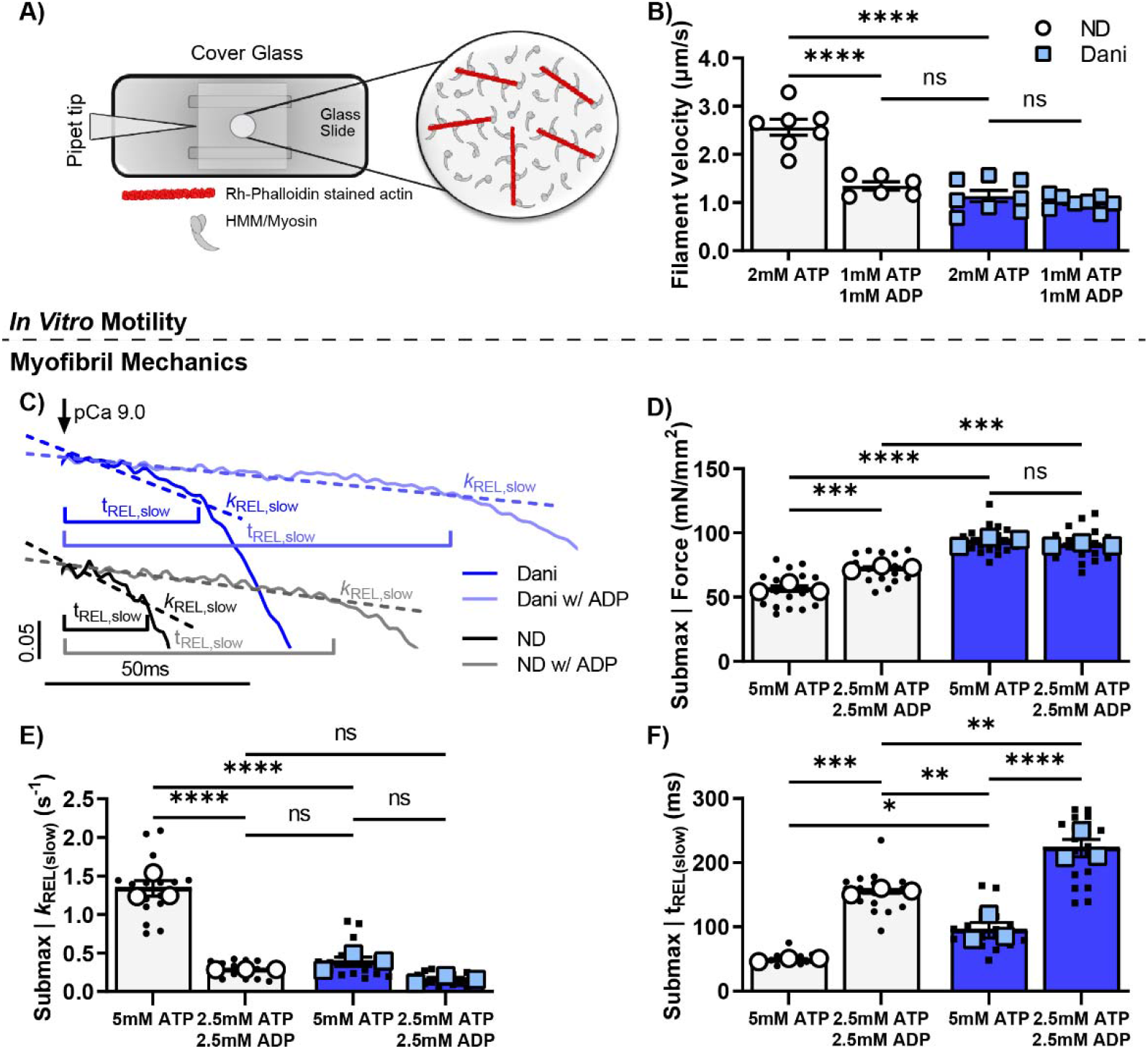
The cross-bridge cycling effects of Danicamtiv are similar to increased ADP. Using the *in vitro* motility assay (A), filament sliding velocity was significantly increased with 0.5 μM Dani treated ATP-bound myosin (left columns). When Dani was applied to myosin with 50% ADP/50%ATP, the filament velocity was unchanged. Myofibril force and activation/relaxation kinetics were measured with/without Dani in presence of ATP or 50%ADP/50% ATP mixture at pCa=5.8. Example tracings are included and are offset for visual clarity (C). The presence of ADP or 1 μM Dani both increased force with the effect not being additive (D). Similarly, ADP slowed *k*_REL(slow)_ to a similar extent as Dani and presence of both did not decrease it further (E). ADP prolonged t_REL(slow)_ to a greater extent than Dani and the effect was cumulative with the combination of the two having the largest effect (F). Data is from 7-8 slides per condition for *in vitro* motility and 18 DMSO or 21 Dani treated myofibrils from 3 biological replicates. *P < 0.05., **P<0.005, ***P<0.001, ****P<0.0001, error bars represent SEM.

**Figure 5.**
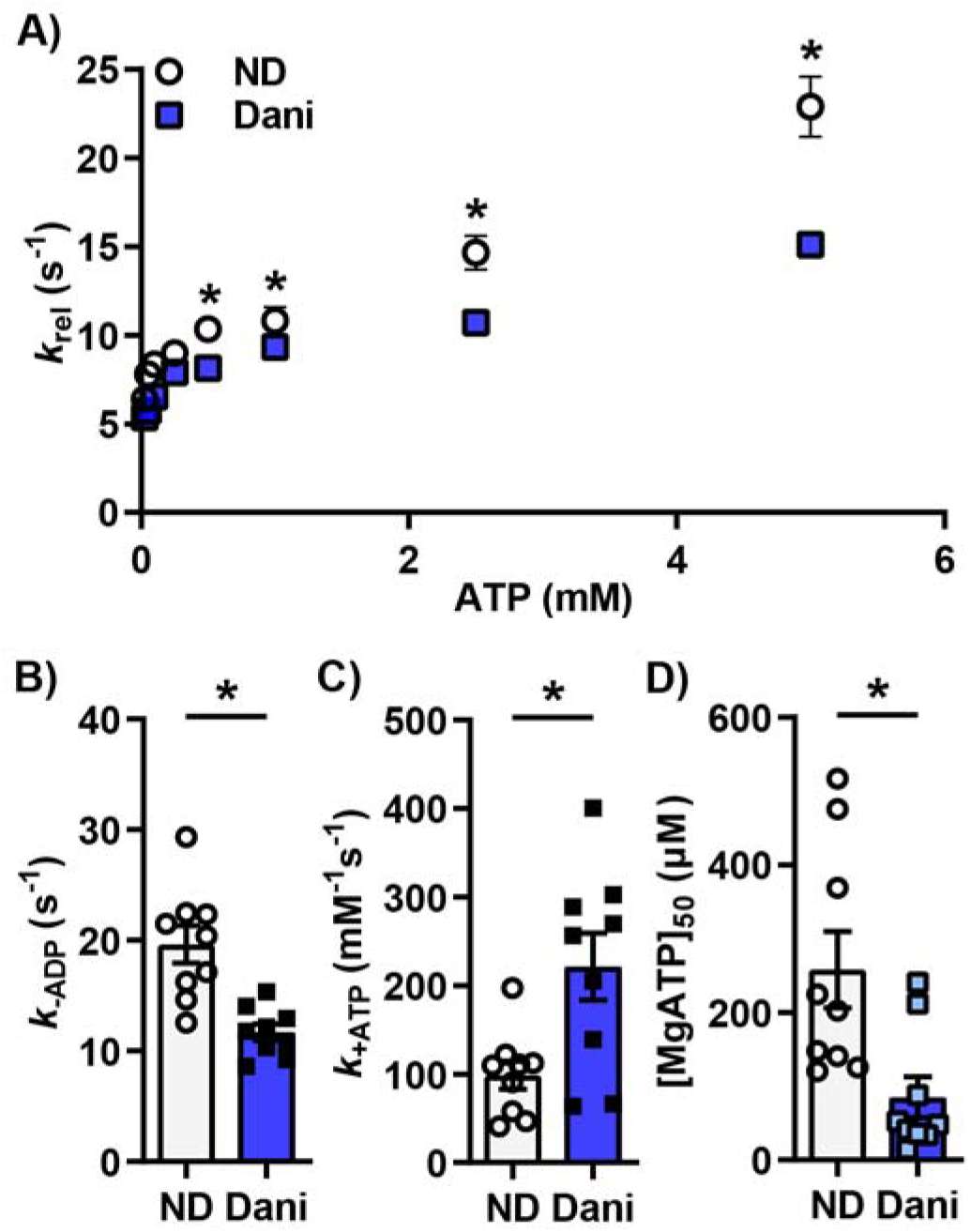
1 μM Danicamtiv increased ATP binding and decreased ADP release in activated porcine cardiac muscle. (A) As expected, the cross-bridge detachment rates (k_rel_) increased with increasing [MgATP] for the untreated and Dani-treated myocardial strips. Treatment with dani decreased *k*_rel_ at [MgATP] grearer than 0.5 mM (B). Fitting the data to Eq. 1 shows that dani decreased ADP release rate (B, *k*_-ADP_) while increasing ATP binding (C, *k*_+ATP_). MgATP concentration at half-maximal detachment rate ([MgATP]50) was lower for the danicamtiv-treated strips (D). Data is from 9 tissue preparations per condition. *P<0.05 vs. ND, error bars represent SEM.

Next, we performed measurements in myofibrils at submaximal calcium (pCa=5.8). Previous studies have shown that under isometric conditions, ADP release is the rate-limiting step in the cross bridge cycle and cardiac muscle relaxation kinetics. Furthermore, by increasing ADP levels, there is an increase in maximal force and calcium sensitivity (36, 38, 39). To ask the specific question of whether Dani directly affects ADP release from myosin, we made our myofibril measurements with Dani in the presence of increased ADP levels. As expected, ADP (Figure 5C-F, S4) increased force (56.6±3.2 vs. 72.6±2.4 mN/mm^2^) and slowed myofibril relaxation rates (*k*_REL(slow)_: 1.340±0.101 vs. 0.286±0.001 s^-1^; *k*_REL(fast)_: 12.81±1.05 vs. 4.34±0.32), while prolonging the duration of the slow phase of relaxation (t_REL(Slow)_: 49.1 ±1.7 vs. 155.2±3.0 ms). Treatment with Dani at submaximal calcium level (pCa=5.8) resulted in a similar increase in force and slowing of activation and relaxation kinetics that was seen at maximal calcium level (Figure 5C-F and Table 1). Combined treatment with Dani and elevated ADP did not further increase force or decrease crossbridge detachment *k*_REL(slow_) compared to Dani and no ADP (Figure 5D-E). This suggests that ADP release is the key step in the cross bridge cycle that is affected by Dani. Interestingly, addition of elevated ADP to Dani treated myofibrils further prolonged thin filament deactivation duration (t_REL(Slow_): 94.9±12.2 vs. 222.5±13.6 ms, P<0.0001), likely due to more strongly-bound cross bridges.

**Table 1.** Numerical values of the mechanical measurements from permeabilized porcine cardiac tissue and myofibrils treated with Danicamtiv. Data is from 19 paired demembranated tissue preparations and 18 DMSO (no drug) or 21 Dani treated myofibrils from 3 biological replicates. Data is reported as mean ± SEM. P values are for comparison of no drug vs. Dani for each parameter using paired two-tailed t-test (demembranated mechanics), Welch’s unpaired two-tailed t-test (myofibrils, pCa 4.0), or two-way ANOVA followed by Tukey’s multiple comparisons test (myofibrils, pCa 5.8).

| **Demembranated Mechanics** | **No drug** | **1 μm Dani** | **P value** |
| --- | --- | --- | --- |
| **Max \| Force (mN/mm<sup>2</sup>)** | 31.80 ± 1.03 | 35.09 ± 1.43 | 0.024 |
| **pCa<sub>50</sub>** | 5.72 ± 0.02 | 5.91 ± 0.05 | 0.033 |
| **Hill Coefficient** | 3.96 ± 0.31 | 2.27 ± 0.29 | 0.0007 |
| **Max \| $k_{tr}$** | 1.91 ± 0.14 | 0.83 ± 0.08 | 0.004 |
| **Myofibrils** | **Maximal Calcium (pCa 4.0)** |  |  |
| **Force (mN/mm<sup>2</sup>)** | 86.07 ± 1.63 | 110.60 ± 2.88 | 0.004 |
| **$k_{ACT}$ (s<sup>-1</sup>)** | 1.24 ± 0.01 | 0.86 ± 0.06 | 0.024 |
| **$k_{REL(slow)}$ (s<sup>-1</sup>)** | 0.77 ± 0.03 | 0.25 ± 0.05 | 0.002 |
| **$t_{REL(slow)}$ (ms)** | 56.5 ± 1.9 | 103.7 ± 9.4 | 0.034 |
| **$k_{REL(fast)}$ (s<sup>-1</sup>)** | 10.64 ± 0.20 | 6.34 ± 0.22 | 0.0001 |
|  | **Submaximal Calcium (pCa 5.8)** |  |  |
| **Force (mN/mm<sup>2</sup>)** | 56.56 ± 3.17 | 93.62 ± 2.17 | <0.0001 |
| **% maximal force** | 65.10 ± 1.47 | 84.66 ± 1.45 | 0.001 |
| **$k_{ACT}$ (s<sup>-1</sup>)** | 0.70 ± 0.01 | 0.53 ± 0.03 | 0.002 |
| **$k_{REL(slow)}$ (s<sup>-1</sup>)** | 1.34 ± 0.10 | 0.39 ± 0.06 | <0.0001 |
| **$t_{REL(slow)}$ (ms)** | 49.1 ± 1.7 | 95.9 ± 12.2 | 0.034 |
| **$k_{REL(fast)}$ (s<sup>-1</sup>)** | 12.81 ± 1.05 | 7.61 ± 0.53 | 0.002 |
| **$k_{ACT}$ (s<sup>-1</sup>) with 50% ADP** | 0.46 ± 0.02 | 0.43 ± 0.02 | 0.781 (ns) |
| **$k_{REL,slow}$ (s<sup>-1</sup>) with 50% ADP** | 0.29 ± 0.01 | 0.16 ± 0.03 | 0.478 (ns) |
| **$t_{REL(slow)}$ (ms) with 50% ADP** | 155.2 ± 3.0 | 222.5 ± 13.6 | 0.004 |
| **$k_{REL(fast)}$ (s<sup>-1</sup>) with 50% ADP** | 4.34 ± 0.32 | 2.39 ± 0.19 | 0.190 (ns) |

**Table 2.** Numerical values of X-ray diffraction reflections of relaxing muscle (pCa 8.0). Values represent mean ± S.E.M. for n= 11 preparations using paired two-tailed t-test analysis.

| **X-ray reflection** | **No drug** | **1 μm Dani** | **P value** |
| --- | --- | --- | --- |
| **d<sub>1,0</sub> (nm)** | 36.62 ± 0.30 | 36.88 ± 0.3 | 0.003 |
| **I<sub>1,1</sub>/I<sub>1,0</sub>** | 0.35 ± 0.02 | 0.47 ± 0.02 | 0.0001 |
| **I<sub>M3</sub> (a.u.)** | 15.6 ± 1.5 | 11.4 ± 1.1 | 0.0101 |
| **I<sub>MLL1</sub> (a.u.)** | 23.2 ± 2.2 | 15.0 ± 1.7 | 0.0002 |
| **I<sub>M6</sub> (a.u.)** | 2.93 ± 0.29 | 2.55 ± 0.31 | 0.114 (ns) |
| **S<sub>M6</sub> (nm)** | 7.209 ± 0.002 | 7.224 ± 0.002 | 0.0008 |

### Nucleotide handling rates were modified in the presence of Danicamtiv

To confirm our myofibril results, we measured changes in nucleotide binding and release rates in presence of Dani. We calculated ATP binding and ADP release rates in porcine cardiac tissue using a step-length protocol in the presence of increasing [MgATP] (Figure S6). Cross bridge detachment rates (*k*_rel_) increased as [MgATP] increased for the untreated and Dani-treated myocardial strips (Figure 4A). *k*_rel_ was slower for the Dani-treated strips at [MgATP] greater than 0.5 mM. Fits to Eq. 1 suggest that slowed cross bridge detachment stems from a combination of slower cross bridge MgADP release rate (*k*_-ADP_) and faster cross bridge MgATP binding rate (*k*_+ATP_) in the presence of Dani (Figure 4B-C). Because the rate limiting step of the crossbridge cycle under load is MgADP dissociation from strongly bound myosin cross bridges in demembranated myocardial strips, it was the slower *k*_-ADP_ that primarily drove the differences in crossbridge kinetics between the untreated and Dani-treated strips. As seen in Figure 4, *k*_+ATP_ was significantly faster than k_-ADP_, and with *k*_+ATP_ being a second-order nucleotide binding process, the [MgATP] needs to be very low for *k*_+ATP_ to affect cross bridge detachment rates. Figure 4D shows that the MgATP concentration at half-maximal detachment rate ([MgATP]_50_) was much lower for the Dani-treated strips. This suggests that Dani-treated preparations required less MgATP to reach their maximal cross bridge detachment rate.

### Danicamtiv recovered abnormal tension in a rodent DCM model

Since Dani is under investigation for treatment of patients with genetic cardiomyopathy, we tested its ability to recover the force deficit in a thin filament DCM mouse model with the characteristic progressive dilation and hypocontractality of human genetic DCM. Permeabilized muscle preparations showed a right-ward shift in the force vs. pCa relationship in the I61Q cTnC DCM model as we have previously reported (12). Additionally, the I61Q cTnC mutant decreased F_max_ and the Hill coefficient. As seen in Figure S5, 1 μm Dani was able to shift the force vs. pCa curves to the left for both control and I61Q cTnC mice with an increase in pCa50 (5.58±0.01 vs. 5.72±0.02 for control, 5.35±0.04 vs. 5.47±0.03 for I61Q cTnC). This increase in pCa50 was significant versus no drug regardless of genotype. Similar to the results in pig cardiac tissue, we found a decrease in the Hill coefficient and maximal *k*_tr_ but with no increase in maximal force (Figure S7 and Table 3). This suggests that effects are independent of myosin isoform.

**Table 3.** Numerical values of mechanical measurements of demembranated cardiac tissue from DCM mice in absence (ND) or presence of 1 μm Danicamtiv (Dani). Values represent mean ± S.E.M. for N = 6-8 animals and <12 preparations per group. * P<0.05 vs. ND, † P<0.005 vs. ND, # P<0.0001 vs. ND using a two-way ANOVA analysis with Šídák’s multiple comparisons test.

| Mechanics Parameter | Control |  | I61Q cTnC |  |
| --- | --- | --- | --- | --- |
|  | ND | Dani | ND | Dani |
| Max \| Force ($\text{mN/mm}^2$) | 64.6 ± 5.8 | 63.1 ± 5.3 | 36.7 ± 4.9 | 41.0 ± 4.2 |
| pCa<sub>50</sub> | 5.61 ± 0.01 | 5.76 ± 0.02<sup>†</sup> | 5.32 ± 0.05 | 5.45 ± 0.03* |
| Hill Coefficient | 3.11 ± 0.09 | 2.15 ± 0.13<sup>#</sup> | 2.45 ± 0.07 | 1.66 ± 0.07<sup>#</sup> |
| Max \| $k_{tr}$ | 28.1 ± 1.4 | 10.8 ± 1.0<sup>#</sup> | 28.2 ± 2.3 | 11.6 ± 1.7<sup>#</sup> |

**Table 4:**
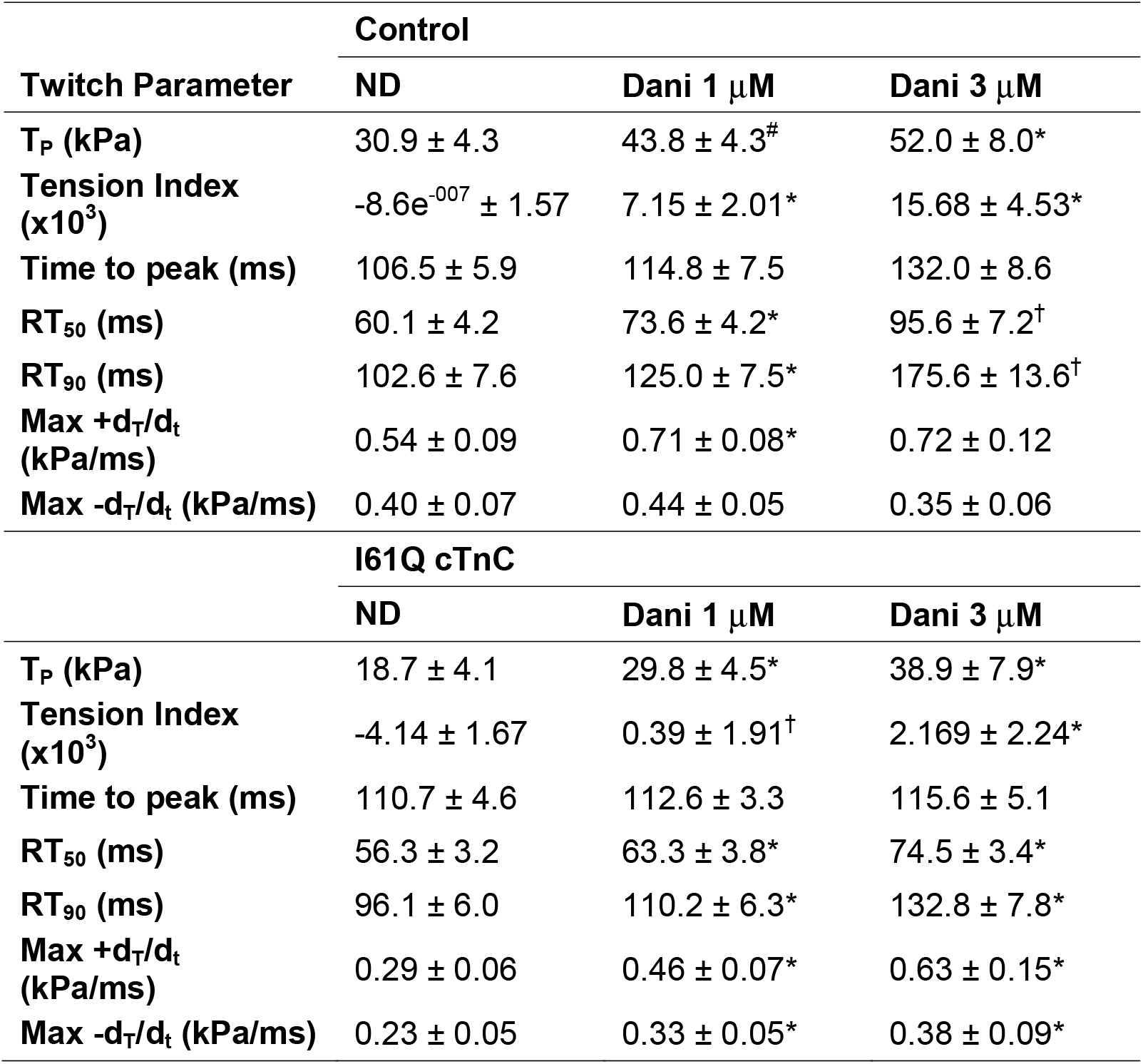
Numerical values of intact trabecula twitch measurements of DCM mice in absence (ND) or presence of 1 or 3 μm Danicamtiv (Dani). Values represent mean ±S.E.M. for 15 control mice and 11 I61Q mice. *P<0.05, † P<0.005, # P<0.0001 vs. ND using a mixed-effect analysis with Dunnett’s multiple comparisons test.

Our intact trabecula measurements showed a decrease in peak tension (T_p_) and tension index in the I61Q cTnC mice (Figure 6) as previously reported (12, 40). In the presence of Dani, T_p_ increased in both control and DCM. In control mice, 1 and 3 μM Dani resulted in ~63% and 131% increase in T_p_, while the increase is ~87% and ~148% respectively in I61Q cTnC trabecula. While the time to peak is unchanged in presence of Dani, time to 50 and 90% relaxation is increased in presence of Dani (Figure S8). We have previously shown that the tension index (TI), the area under the twitch curve that is subtracted from a healthy twitch, is predictive of cardiomyopathy phenotype (12, 13). While TI increased with Dani in a dose dependent manner, the increase was seen more in the control mice. This is mostly due to the more pronounced relaxation change in the control tissue. Using mixed-effects analysis, the interaction between genotype and Dani is statistically significant, suggesting a different treatment response in control versus I61Q cTnC tissue. These results suggest that the underlying mechanism of cardiomyopathy might need to be considered when dosing a medication such as Danicamtiv.

**Figure 6.**
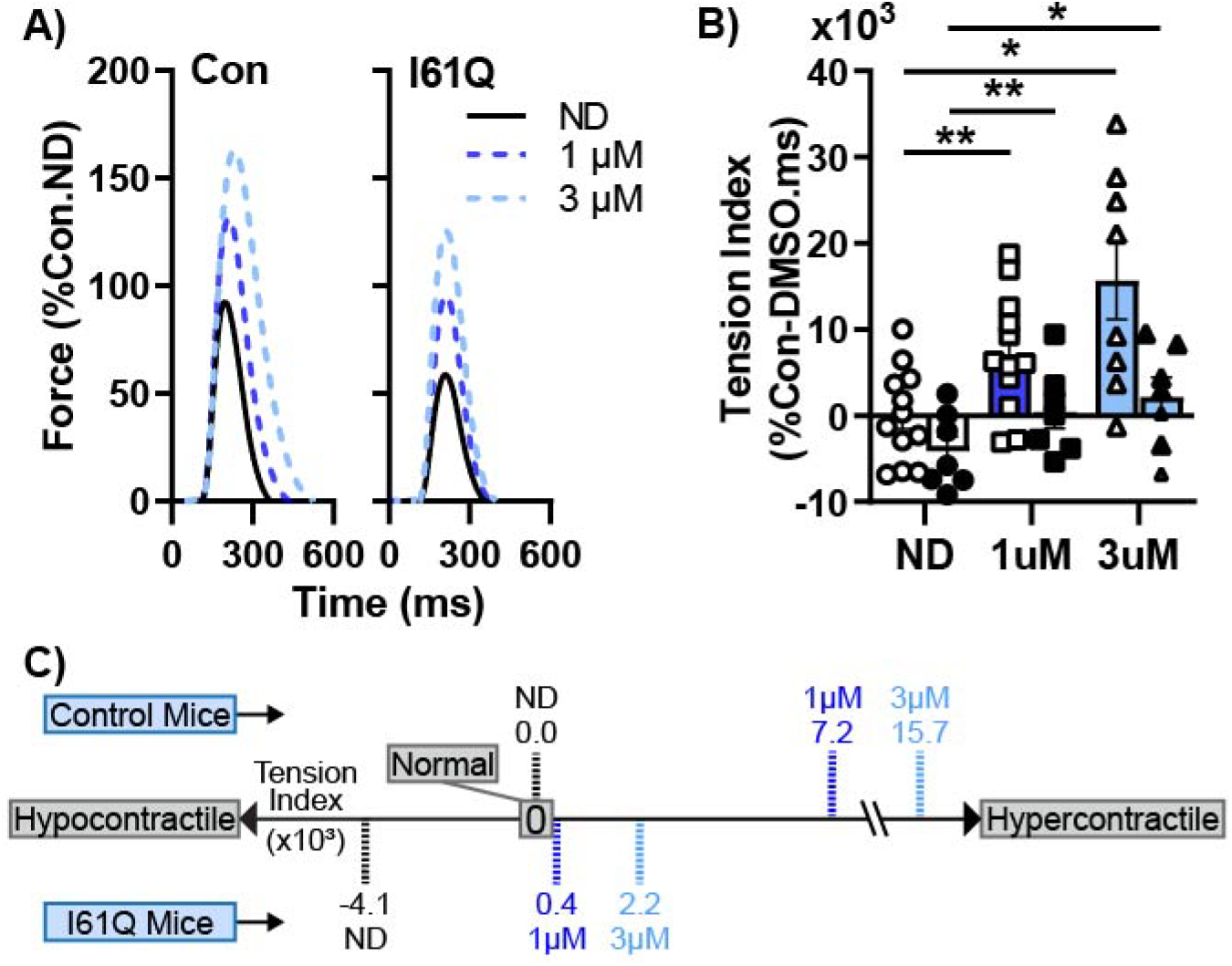
Dani mitigated the contractile abnormalities in a rodent DCM model. (A) Average twitch force-time traces (in % Control-ND T_p_) of intact papillary muscle trabeculae from control (left) and I61Q cTnC (right) mice stimulated at 1 Hz. Dani increased Tp in control and I61Q cTnC twitches in a dose dependent manner. (B) While Dani increased tension index in both control (open symbols) and I61Q cTnC (closed symbols), the control mice responded more favorably (C). Data is from 15 control mice and 11 I61Q mice. *P<0.05 and **P< 0.01 vs. ND, error bars represent SEM.

## Discussion

We set out to perform a first detailed analysis of the mechanism of action of the novel myosin activator Danicamtiv. Our studies can provide a methodological framework to evaluate how small molecules modulate specific aspects of the cross bridge cycle that affect cardiac myofibril activation and relaxation. The data can also help identify subsets of patients that may derive the most benefit from use of this myosin activator. For example, there could be a different response in sarcomeric versus non-sarcomeric DCM or thin filament versus thick filament variants. Genetic variants are being recognized as an increasing cause of heart failure and dilated cardiomyopathy. Myosin activators may provide a treatment that addresses the causative hypocontractile phenotype. This motivated a current clinical trial of patients with genetic DCM for treatment with Dani. However, there is lack of data regarding the utility of this medication in a genetic cardiomyopathy preclinical model and if patients with different causative genes may derive differential benefits. Using a rodent model of genetic DCM with reduced thin filament activation, we were able to assess the ability of Dani to correct the hypocontractility in this model and compare its effectiveness to treatment in control mice.

There are several key findings that emerged from this study. First, we showed that treatment with Dani restructured the thick filament of resting muscle to a more ON configuration, where myosin heads are positioned closer to actin. This increased the number of myosin heads available to contribute to contraction, a result that is consistent with more myosin motors binding at lower calcium concentrations. It also suggests reduced reliance on calcium mediated cooperativity of myosin binding, hence an increase in calcium sensitivity with a decreased Hill coefficient. Second, we showed that Dani slowed cross bridge turnover and subsequently decreased rate of myofibril relaxation and prolonged thin filament deactivation. This prolonged relaxation manifested in intact cardiac muscle, which had slower twitch relaxation kinetics. Third, we provided evidence to show that the decrease in myosin ADP release rate is the mechanism for decreased cross bridge turnover and slower relaxation. Fourth, we showed that Dani improved the tissue level hypocontractility in a genetic DCM model.

### Danicamtiv restructures myosin in an analogous manner to other myosin activators

Myosin activation is a novel way to treat heart failure. Omecamtiv mecarbil (OM), the first in class myosin activator, recently completed a phase 3 clinical trial for treatment of systolic heart failure patients (9). Studies using fluorescent probes have demonstrated that OM similarly stabilizes the ON state of the thick filament resulting in more myosin motors available for binding to the thin filament (10). Our x-ray diffraction studies also showed partial thick filament activation with Dani under resting conditions. This activation under resting conditions can be expected to result in more myosin motors binding at lower calcium concentrations with less reliance on cooperativity. Results of our sinusoidal length-perturbation analysis further support that Dani increases cross bridge binding and promotes an increase in the ratio of ON to OFF cross bridge populations. Similar changes in myosin structure under resting conditions are induced by 2-deoxy adenosine triphosphate (dATP), a nucleotide known to increase force and calcium sensitivity in rodent, dog, and human cardiac myocardium (17, 41, 42), correlate well with active force (26, 43).

### Danicamtiv inhibits cross bridge cycling rate, prolonging thin filament deactivation and relaxation in myofibrils

The myofibril kinetics measurements suggest that Dani prolongs thin filament activation, as indicated by the longer t_REL(slow)_ and that this is due to a decreased rate of cross bridge detachment as measured by *k*_REL(slow)_. A similar mechanism of action involving prolonged actomyosin attachment resulting in cooperative thin filament activation has been proposed as a mechanism for the myosin activator OM**Error! Bookmark not defined.**(7, 23, 44). The inhibitor-like mechanism is further supported by decreased ATP turnover in HMM with increasing concentrations of Dani (Figure S7). The duration of the slow phase of relaxation (t_REL(slow)_) is known to be affected by changes in troponin complex calcium affinity (20) so we measured the effect of Dani on troponin function. Dani did not change Ca^2+^ binding to cardiac troponin C (cTnC) as measured by steady state fluorescence spectroscopy using site specific labeling at C84 of cTnC with fluorescence probe IANBD (Supplemental Figure S9). Therefore, the decreased cross bridge detachment rate is likely explained by the decreased ADP release rate, and this is supported by myofibril relaxation measurements in the presence of elevated ADP, which mimic relaxation kinetic changes seen with Dani under conditions of load.

### Danicamtiv corrects abnormal contraction in a genetic DCM mouse model

Using a rodent model of sarcomeric genetic DCM with decreased thin filament activation, we demonstrated that Dani could normalize the decreased calcium sensitivity of contraction in I61Q cTnC mice. At the lowest calcium levels of our F-pCa curve, there is full recovery of the force deficit to levels of control myocardium. However, the maximum force is still significantly lower in the Dani-treated I61Q cTnC trabeculae. The decreased *k*_tr_ in mice suggested that slowing of cross bridge kinetics was present in hearts with both MYH6 (mice) and MYH7 (pig). Intact trabecula experiments demonstrated that tension increased in both control and I61Q cTnC mice after treatment with Dani. Control mice consistently achieved a higher peak force and tension index, a summative measure encompassing peak tension and kinetics of both activation and relaxation, compared to I61Q. Control mice also expressed greater sensitivity to Dani treatment with significantly greater time to 50% and 90% relaxation compared to I61Q. An explanation for the difference in relaxation is that Dani normalizes the duration of thin filament deactivation in the I61Q cTnC which are abnormal without treatment. However, in the control mice, the thin filament stays on much longer, resulting in a much longer relaxation. The different sensitivity to Dani treatment due to genotype can be summarized by a statistically significant interaction between genotype and treatment for the tension index.

Our study has some limitations. Our mouse model of genetic DCM is based on an engineered mutation in cTnC and not a known disease-causing mutation. It is also worth noting that MYH6 is the dominant form of cardiac myosin in mice, while humans express the MYH7 variant. It is possible that our genetic DCM results could be different in organisms with MYH7 as the dominant myosin. Intact twitch measurements were only performed in mice, which expresses the fast form of myosin. Our studies have not assessed the relative affinities of Dani for different cardiac myosin isoforms, however we found comparable results in mouse and pig, which have different predominant isoforms.

In conclusion, we use a variety of different tools to demonstrate the mechanisms of how Danicamtiv works as a myosin activator to increase force and calcium sensitivity. The initial discovery of Dani was based on increased myofibril ATPase activity. Our studies support a more complex process than a simple increase in myosin cycling rate. The inhibition of ADP release leads to slower relaxation kinetics of myofibrils and consequently slower relaxation in intact tissue. This widening of the twitch duration contributes to the increase in tension index. In addition, myosin heads are more primed to interact with actin at low levels of calcium, resulting in increased calcium sensitivity and higher contractile forces at levels of calcium the sarcomere experiences during the cardiac cycle. This increase in force production also contributed to the increased tension index. Together, all these mechanisms lead to increased overall function in cardiac tissue from rodent genetic DCM hearts. Future studies will need to look at a diverse group of variants and extend the studies to human patients with genetic DCM.

## Materials and Methods

### Animal use and ethics

All experiments followed protocols approved by both the University of Washington and the Illinois Institute of Technology Institutional Animal Care and Use Committees according to the “Guide for the Care and Use of Laboratory Animals” (National Research Council, 2011). Farm pig hearts were obtained immediately after animal was euthanized and rinsed in cold oxygenated Tyrode’ buffer.

### Dilated cardiomyopathy mouse model

We used a previously published rodent genetic DCM model. The I61Q cTnC is an engineered cTnC variant that is known to desensitizes the myofilament to calcium via altering cTnC calcium binding (20, 45). Transgenic mice show approximately 45% replacement of the normal cTnC with I61Q cTnC and progressive ventricular dilation and reduced contractility as they age (12). Experimental mice were between 3-5 months of age at time of euthanasia.

### Excision of mouse hearts

Hearts were rapidly excised via thoracotomy and immediately rinsed in an oxygenated (95% O_2_, 5% CO_2_) modified Krebs buffer containing (in mM) 118.5 NaCl, 5 KCl, 1.2 MgSo_4_, 2 NaH_2_PO_4_, 25 NaHCO_3_, 1.8 CaCl_2_, and 10 glucose. The solution was bubbled with 95% O_2_/5% CO_2_ for a minimum of 10 minutes to bring pH to ~7.2 before addition of CaCl_2_. Hearts were then perfused, and ventricles splayed open in oxygenated modified Krebs with 0.1 CaCl_2_ and 20 2,3-butanedione 2-monoxime (BDM) to inhibit contraction and minimize damage during tissue dissection.

### Demembranated tissue mechanics

#### Isometric force and rate of tension redevelopment

Frozen porcine left ventricular tissue or freshly excised mouse hearts were permeabilized in 50:50 glycerol relaxing solution containing (in mM) 100 KCl, 10 MOPS, 5 K_2_EGTA, 9 MgCl_2_ and 5 Na_2_ATP (adjusted to pH = 7 with KOH), 1% (by vol) Triton X-100, 1% protease inhibitor (sigma P8340), and 50% (by vol) glycerol at 4°C overnight. After permeabilization, the solution was changed to the same 50:50 glycerol relaxing solution without Triton X-100 for storage up to one week at −20°C. The mouse tissue was used fresh. Pig tissue data includes that from both fresh and previously frozen tissues with no significant difference. Permeabilized trabeculae/papillary muscles (mouse) or thin strips (pig) were dissected and mounted between a force transducer (Aurora Scientific, model 400A) and a motor (Aurora Scientific, model 315C) using aluminum T-clips (Aurora Scientific) as previously described (13). Sarcomere length (SL) was set to ~2.3 μm for the experiments. Experiments were conducted in physiological solution (pH 7.0) at 15°C (mouse) or 21°C (pig) containing a range of pCa (= −log_10_[Ca^2+^]) from 9.0 to 4.0 with and without 1μM Dani. Tissue was allowed to reach stead-state force (F) at each pCa. F-pCa curves were collected and analyzed with custom code using LabView software and fit to the Hill equation to calculate the pCa_50_, the pCa at half-maximal force, and the Hill coefficient (nH), a measure of the cooperativity of force. The rate of tension redevelopment (*k*_tr_; following a 15% rapid release-restretch transient) was calculated from the half time of force recovery. High frequency stiffness (HFS) was measured by applying a 1000 Hz sinusoidal length change (±0.5% muscle length (ML)) and was calculated from the ratio of peaks of the Fourier transforms of the force and ML signals.

#### Phosphate inhibition

Experiments assessing the effects of inorganic phosphate (Pi) were done at 21°C in equivalent physiological solutions as above made at pCa 9 and pCa 4, where Pi was incorporated into the recipe for a final [Pi] of 0, 1, 3, and 10 mM Pi.

#### Quick stretch analysis

Solution recipes were calculated and prepared as follows ^1^. Relaxing solution (in mM): pCa 8.0, 5 EGTA, 5 MgATP, 1 Mg^2+^, 0.3 Pi, 35 phosphocreatine, 300 U/mL creatine kinase (CK), 200 ionic strength, 3% dextran T-500 (w/v), pH 7.0. Activating solution was the same as relaxing solution but adjusted to a free [Ca^2+^] of pCa 4.8. Dissecting solution (in mM): 50 BES, 30.83 K propionate, 10 Na azide, 20 EGTA, 6.29 MgCl2, 6.09 ATP, 1 DTT, 20 BDM, 50 Leupeptin, 275 Pefabloc, and 1 E-64, pH 7.0. Permeabilizing (skinning) solution: Dissecting solution with 1% Triton X-100 wt/vol and 50% glycerol wt/vol. Storage solution: Dissecting solution with 50% glycerol wt/vol.

Permeabilized porcine myocardial strips were trimmed to ~180 μm in diameter and ~700 μm in length. Aluminum T-clips were attached to the end of each strip and then mounted between a piezoelectric motor (P841.40, Physik Instrumente, Auburn, MA) and a strain gauge (AE801, Kronex, Walnut Creek, CA). They were lowered into a 30 μL droplet of relaxing solution maintained at 28°C and set at 2.3 μm sarcomere length by stretching the muscle roughly 25% of muscle length from taut. Measurements were made in activating and rigor solutions in the presence and absence of 1μM Danicamtiv under experimental conditions described below.

Stress (=force per cross-sectional area) responses were recorded following a steplength change of 0.5% muscle length to assess cross-bridge kinetics as a function of [MgATP]. The stress response was fit to a dual exponential function to characterize the rate of stress release (*k*_rel_) associated with the cross-bridge detachment rate as [MgATP] varied (Figure S6). Given that strongly bound cross-bridge attachment events occupy the myosin-MgADP state and rigor state of the cross-bridge cycle as [MgATP] was titrated from 5 mM towards 0 mM, krel as a function of [MgATP] can be described by Eq. 1. As [MgATP] decreases, cross-bridge detachment slows due to slower MgATP binding, which increases the amount of time a cross-bridge spends in the rigor state.

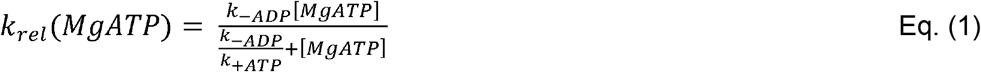

This *k*_rel_ vs. MgATP relationship was fit to Eq. 1 for each individual fiber to estimate the cross-bridge rates of MgADP release (k_-ADP_) and MgATP binding (k_+ATP_). The ratio of *k*_-ADP_/*k*_+ATP_ represents the MgATP concentration at half-maximal detachment rate [MgATP]_50_.

#### Sinusoidal length-perturbation analysis

Sinusoidal length perturbations of 0.125% myocardial strip length (clip-to-clip) were applied at 48 discreet frequencies from 0.125 to 250 Hz to measure the complex modulus as a function of frequency (46–48). The complex modulus represents viscoelastic myocardial stiffness, which arises from the change in stress divided by the change in muscle length that is in-phase (elastic modulus) and out-of-phase (viscous modulus) with the sinusoidal length change at each frequency.

The characteristics of the elastic and viscous moduli responses over the measured frequency range provide a signature of cross bridge binding and cycling kinetics. Vertical shifts in the elastic and viscous moduli are useful for assessing changes in the number of bound cross-bridges between experimental conditions. Frequency-dependent (horizontal) shifts in the moduli are useful for assessing changes in the work-producing and work absorbing characteristics of the myocardium that arise from differences in force generating cross-bridge kinetics.

### Myofibril mechanics

Myofibril preparations were isolated from membranate frozen pig cardiac tissue as previously described (17). Briefly, cardiac tissues were rinsed in Rigor Buffer containing (in mM), 50 Tris, 100 KCl, 2 MgCl_2_, 1 EGTA, pH 7.0, and 1x protease inhibitor (Sigma-Aldrich, St. Louis, MO). Homogenized tissue generated myofibrils using 2 x 30 sec pulses of (VWR Radnor, PA) at low speed and stored at 4 °C to be used for up to two days. Myofibril activation and relaxation measurements were performed on a custom set up as previously described (17, 18). Briefly, myofibrils were mounted between two glass needles; one which acted as a force transducer and the other as an inflexible motor arm mount. A dual diode system measures force based on needle deflection, with force transducer needle stiffness measured at 7.98 mm/μN. A double-barreled glass pipette delivered relaxing (pCa = 9.0) and activating (pCa = 5.8 or 4.0) solutions to the mounted myofibril. Activation and relaxation data were collected at 21 °C and fitted with 50% time to peak, linear interpolation, and 50% time to decay as previously described (17).

### Intact twitch assay

Unbranched, intact trabeculae or papillary muscles were dissected from the right ventricular wall and mounted between a force transduce (Cambridge Technology, Inc., model 400A) and a rigid post. Each end of the trabecula was sutured to custom arms, made from 22-gauge needles, attached to the post and the force transducer. The tissue was then submerged in a custom experimental chamber that was continuously perfused with oxygenated modified Krebs buffer (1.8 mM CaCl_2_) at 33° C. Field stimulation with custom platinum plate electrodes with oscillating polarity triggered muscle twitches. Tissues were lengthened to just taut and paced at 0.5 Hz for ~20 minutes to allow BDM to wash out and tissue to equilibrate. Optimal length was set by stretching the tissue until peak twitch height no longer increased. Continuous twitch tension traces were recorded using custom LabView software at a sampling rate of 1 kHz and were analyzed with custom code written using MATLAB software (version 2021a, The MathWorks).

### In vitro motility assay

Myosin was purified from pig left ventricular samples and then heavy meromyosin (HMM) was prepared by digestion of myosin with 0.05 mg/ml chymotrypsin as previously described (17). Isolated HMM (0.37 mg/ml) was rapidly frozen and stored in Buffer D solution (in mM): 2 MgCl2, 5 EGTA, 5 DTT, Imidazole, 0.2 PMSF, pH 7.4 with 1% sucrose and 1% protease inhibitor at −80 °C. *In vitro* motility assays were performed at 30 °C using unregulated Rhodamine Phalloidin labeled F-actin in the presence of 2 mM ATP and DMSO or 0.5 μM Dani as previously described.(17, 18) In a subset of experiments, the nucleotide composition was changed to 1 mM ATP and 1 mM ADP. Custom-built software was used to analyze images of the moving filaments as in our previous publications (17, 18).

### Stopped flow assay

Stopped-flow experiments were conducted on a HiTech TgK Scientific DX stopped-flow spectrometer as previously described (49). All experiments were excited with 365 nm wavelength light and emission was measured by a PMT through a DD400 long pass filter, unless otherwise stated. HMM was prepared as described in the in vitro motility section, above. All experiments were run in Rigor Buffer as described in the myofibril section, above.

#### ATP binding

HMM and mantATP (fluorescently-labeled ATP; Jena Bioscience) were rapidly mixed inside the stopped flow observation chamber to a final concentration of 100 nM HMM and mantATP ranging from 0.5 – 4 μM. Change in fluorescence was fit to a single exponential growth curve to derive the k_obs_ at each [mantATP]. This yielded a plot of k_obs_ vs [mantATP], with the slope defining the apparent 2^nd^ order rate constant of ATP binding. This was calculated for 0, 1, 3, and 10 μM danicamtiv.

#### ATP turnover

HMM was incubated with mantATP for 1 minute to allow binding then rapidly mixed with unlabeled ATP in the stopped flow observation chamber to final concentrations of 250 nM HMM, 2 μM mantATP, and 125 μM unlabeled ATP. Change in fluorescence was fit to a single exponential decay and the rate of decay was defined as the rate constant for ATP turnover. This was calculated for 0, 1, 3, 10, and 15 μM danicamtiv.

### X-ray diffraction

Wild type Yucatan mini-pig hearts were provided by Exemplar Genetics LLC. Humane euthanasia and tissue collection procedures were approved by the Institutional Animal Care and Use Committees at Exemplar Genetics. Permeabilized tissues were prepared as described previously (50, 51). Briefly, frozen left ventricle wall tissues (about 0.5 −1 cm^3^) was defrosted at room temperature in skinning solution (in mM: 91 K+-propionate, 3.5 MgCl_2_, 0.16 CaCl_2_, 7 EGTA, 2.5 Na_2_ATP, 15 Creatine phosphate, 20 Imidazole, 30 BDM, 1% Triton-X100 and 3% Dextran at pH 7) before dissecting into smaller strips (~5 −10 mm long and 1-2 mm wide). The tissues were permeabilized at room temperature for 3 hours before further dissected into preparations 4 mm long with a diameter of ~200 μm before attaching aluminum T-clips to both ends and stored in pCa8 solution with 3% dextran on ice for the day’s experiments.

X-ray diffraction experiments were performed at the BioCAT beamline 18ID at the Advanced Photon Source, Argonne National Laboratory (52). The X-ray beam energy was set to 12 keV (0.1033 nm wavelength) at an incident flux of ~5 ×10^12^ photons per second. The specimen to detector distance was ~ 3 m. Skinned muscle preparations were mounted in a custom mechanics rig allowing for simultaneous X-ray diffraction and force measurements as monitored by an ASI 610A data acquisition and control system (Aurora Scientific). The muscle was incubated in a customized chamber whose bottom is attached to a heat exchanger, so the solution was kept between 28 °C to 30 °C. The muscles were stretched to a sarcomere length of 2.3 μm using micromanipulators attached to the hooks while monitoring light diffraction patterns from a helium-neon laser (633 nm) on a screen. X-ray fiber diffraction patterns were collected in pCa 8 solution in the absence or presence of 50 μM of danicamtiv on a MarCCD 165 detector (Rayonix Inc., Evanston IL) with a 1 s exposure time. To minimize radiation damage, the muscle samples were oscillated along their horizontal axes at a velocity of 1 - 2 mm/s. The irradiated areas were moved vertically after each exposure to avoid overlapping X-ray exposures. X-ray diffraction patterns were analyzed using the MuscleX software package developed at BioCAT (53). The equatorial reflections were analyzed by the Equator module in the MuscleX software package as described previously (54). X-ray patterns were subsequently quadrant folded and background subtracted to improve signal to noise ratio for further analysis using the Quadrant Fold module of the MuscleX program suite. The meridional and layer line reflections were measured using the Projection Traces module of MuscleX program suite as described (55, 56). Three to four patterns were collected under each condition and the X-ray reflection data extracted from these patterns were averaged.

### Statistical Analysis

We used GraphPad Prism 9 for data and statistical analysis. The data is presented as mean with error bars representing the standard error of the mean. For porcine demembranated tissue mechanics, we used a mixed-effect model with Šídák’s multiple comparisons test for pCa curves and paired two-tailed t-test for the remaining measures. For myofibrils, we used Welch’s unpaired two-tailed t-test (pCa 4.0), or two-way ANOVA followed by Tukey’s multiple comparisons test (pCa 5.8 ± ADP). For X-ray diffraction, we used paired two-tailed t-tests. For *in vitro* motility, we used two-ANOVA with Šídák’s multiple comparisons. For quick step and sinusoidal length-perturbation assays, statistical analyses were performed using SPSS (IBM Statistics, Chicago, IL). We used nested linear mixed models incorporating two effects (treatment and [ADP]) to assess differences of k_rel_ at each [ADP] by treatment group. We used nested linear mixed model analyses to link data from the same hearts, with hearts being a random effect, to optimize statistical power. We used unpaired Student’s t-test to assess differences of nucleotide handling rates. Post hoc analyses were performed using Fisher’s least significant difference test, where p < 0.05 were considered significant. For murine intact mechanics, we used mixed-effect analysis with Dunnett’s multiple comparisons. For murine demembranated tissue mechanics, we used two-way ANOVA analysis with Šídák’s multiple comparisons.

## Acknowledgments

This work was supported by NIH Grants R01HL157169 (FMH), R01HL128368 (MR), RM1GM131981 (MR), R01HL142624 (JD), R01HL1656450 (BCWT), and American Heart Association Collaborative Sciences Award (FMH and JD). The authors would like to acknowledge Mr. Darron Marzolf who provided fresh pig tissue for experimentation. This project used resources from the UW (University of Washington) Center for Translational Muscle Research (CTMR). The UW CTMR is supported by the National Institute of Arthritis and Musculoskeletal and Skin Diseases of the National Institutes of Health under Award Number P30AR074990. This research used resources of the Advanced Photon Source, a US Department of Energy (DOE) Office of Science User Facility operated for the DOE Office of Science by the Argonne National Laboratory under Contract DE-AC02-06CH11357. BioCAT is supported by grant P30 GM138395 from the National Institute of General Medical Sciences of the National Institutes of Health. We acknowledge Dr. David Mack for providing access to BioRENDER.

## Supplemental Data

**Figure S1.**
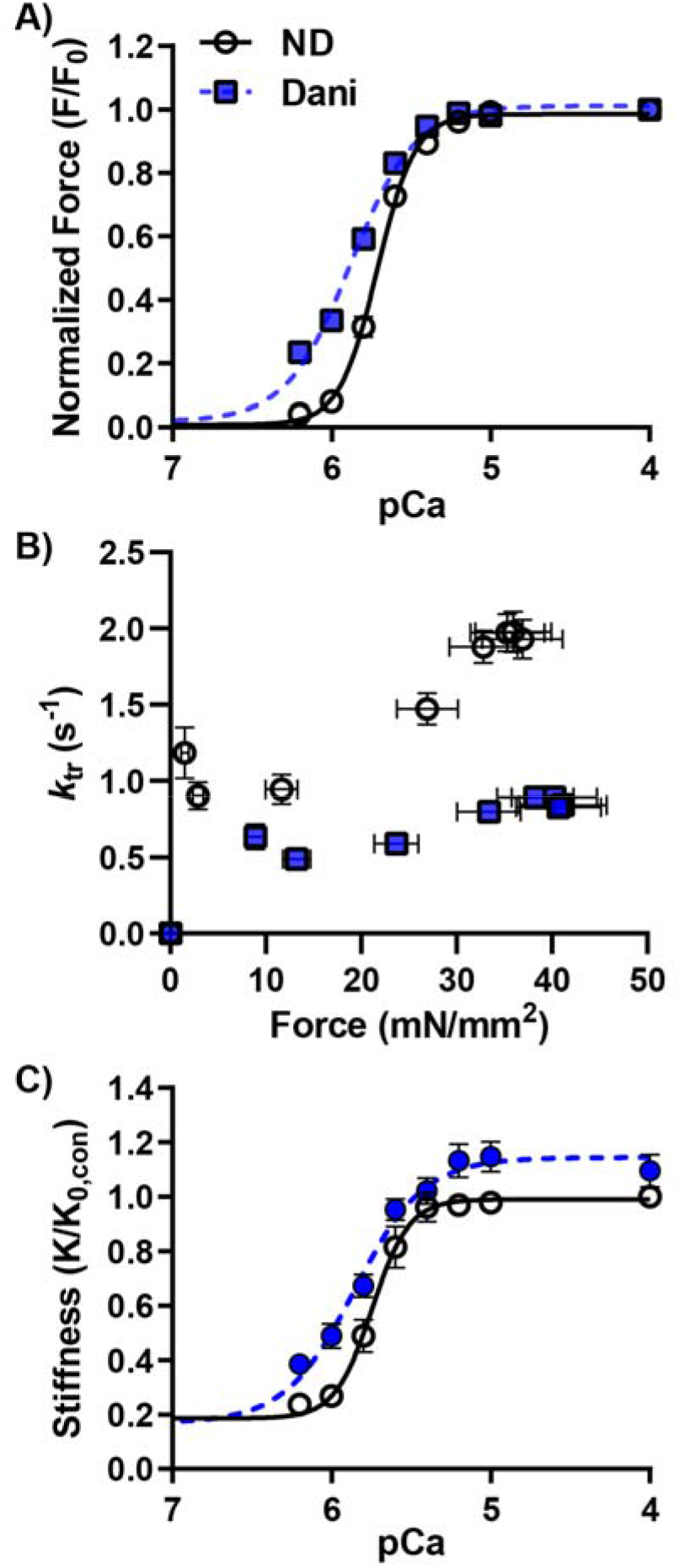
Permeabilized mechanics in porcine cardiac tissue with 1μM Danicamtiv. Force, normalized to maximal force in each condition, versus pCa showed a left shift of the curve with a decrease in the hill efficient (A). Dani reduced force redevelopment rate constants (*k*_tr_) at comparable force measurements. Stiffness, normalized to maximal stiffness without drug, increased proportionally to force(C), suggesting an increase in the number of cross bridges bound with no change in force per cross bridge.

**Figure S2.**
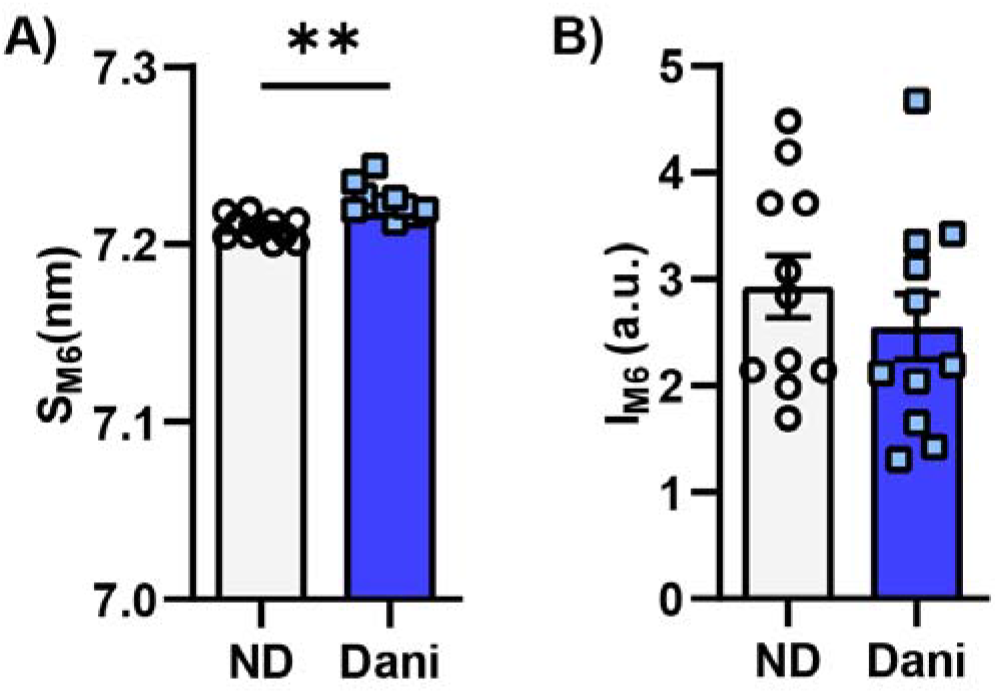
Dani altered the thick filament backbone structure. 50 μM Dani increased spacing of the M6 reflections (A) without changing its intensity (B).

**Figure S3.**
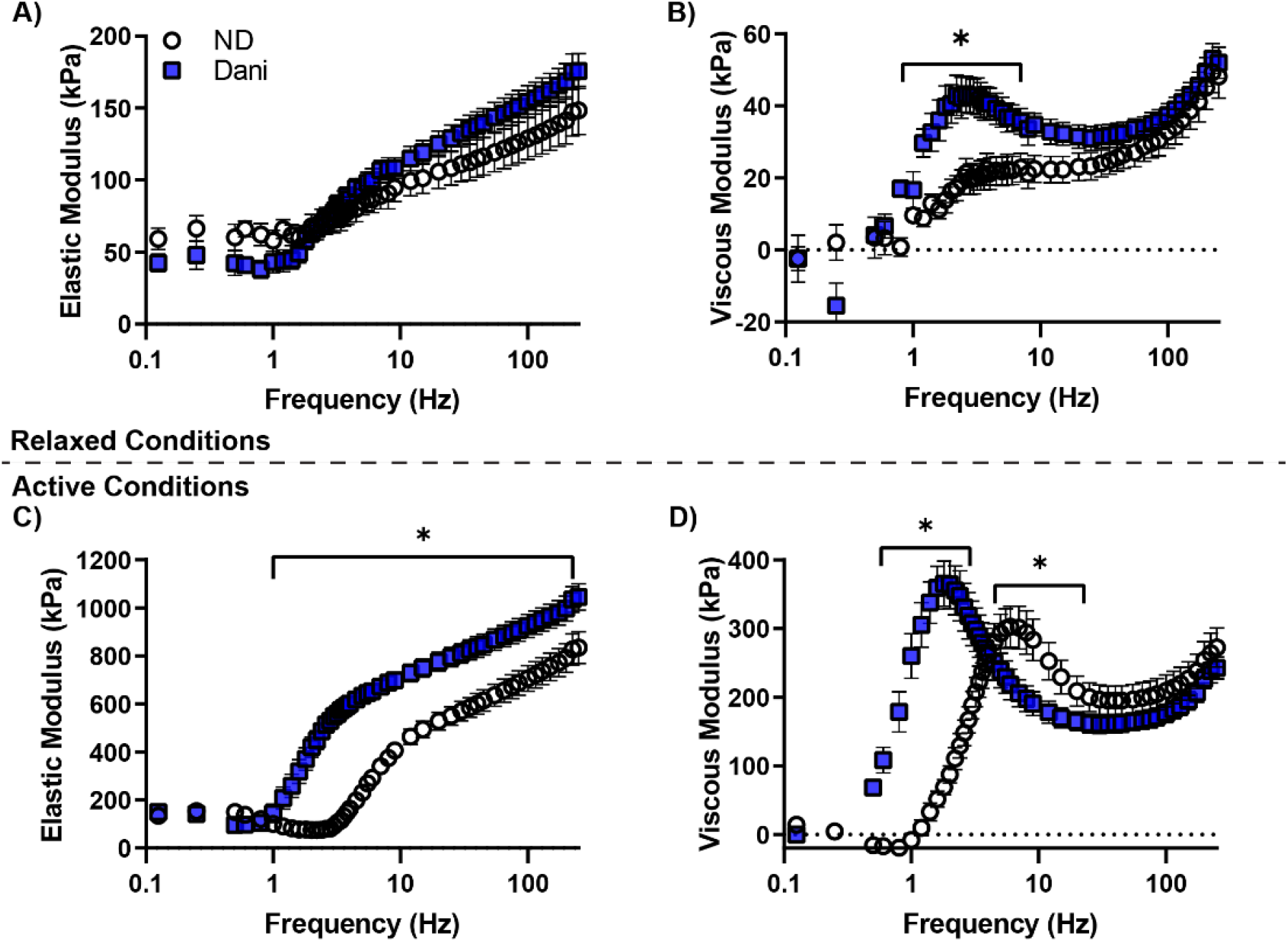
Passive and active elastic and viscous moduli were modified in the presence of 1 μM Danicamtiv. In relaxed muscle, Dani did not alter the elastic moduli (A). Under similar conditions, Dani increased the viscous moduli at a sub-set of oscillatory frequencies, suggesting greater cross bridge binding or cross bridge activity(B). In activated muscle, both elastic (C) and viscous (D) moduli values were greater in the Danicamtiv-treated preparations across a wide range of frequencies, suggested greater cross bridge binding.

**Figure S4.**
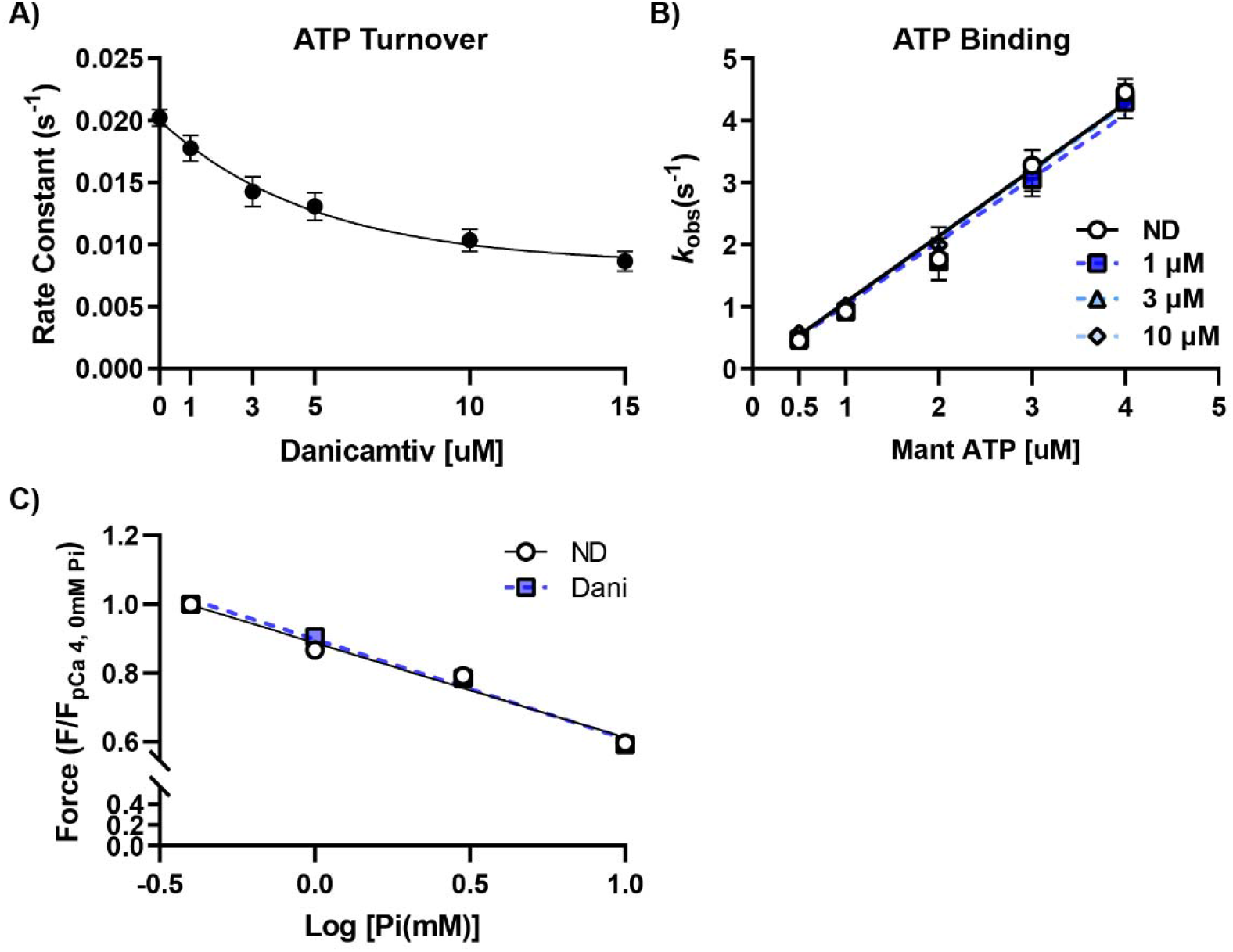
Danicamtiv slowed cross bridge turnover without altering Pi release. Increasing dose of Danicamtiv inhibited the rate constant for ATP turnover in porcine cardiac heavy meromyosin (A). Slope of *k*_obs_ vs [mantATP], which is the 2nd order rate constant of ATP binding, was unchanged after treatment with 1, 3, and 10 μM danicamtiv (B). Treatment with Dani did not alter the linear relationship between relative force and Log [Pi] in permeabilized mechanics in porcine cardiac tissue.

**Figure S5.**
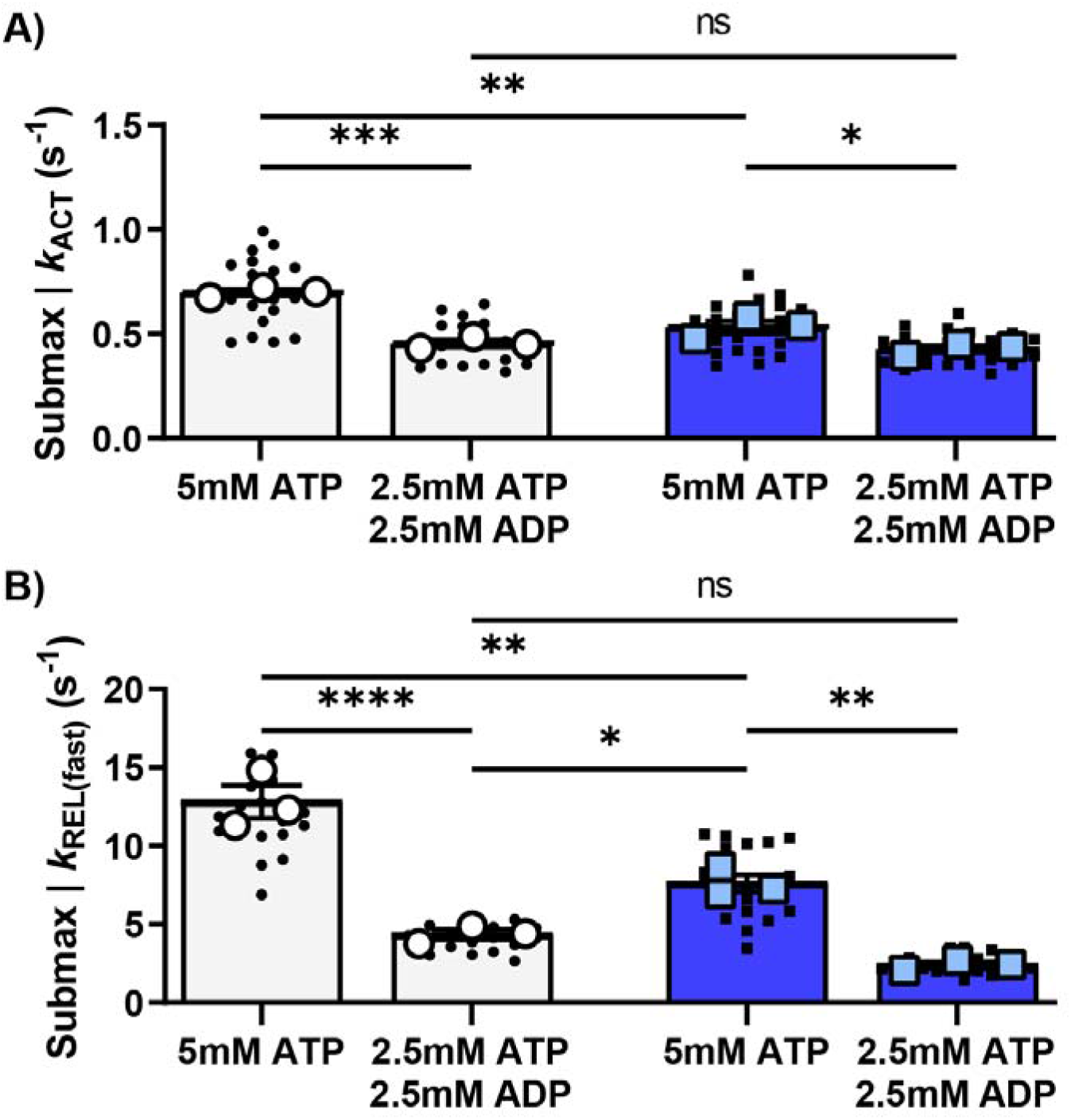
1 μM Danicamtiv and ADP altered myofibril activation and relaxation rates. Dani and ADP decreased exponential activation (A) and relaxation (B) kinetics in porcine myofibrils. The combination of ADP and Dani decreased these rates further.

**Figure S6.**
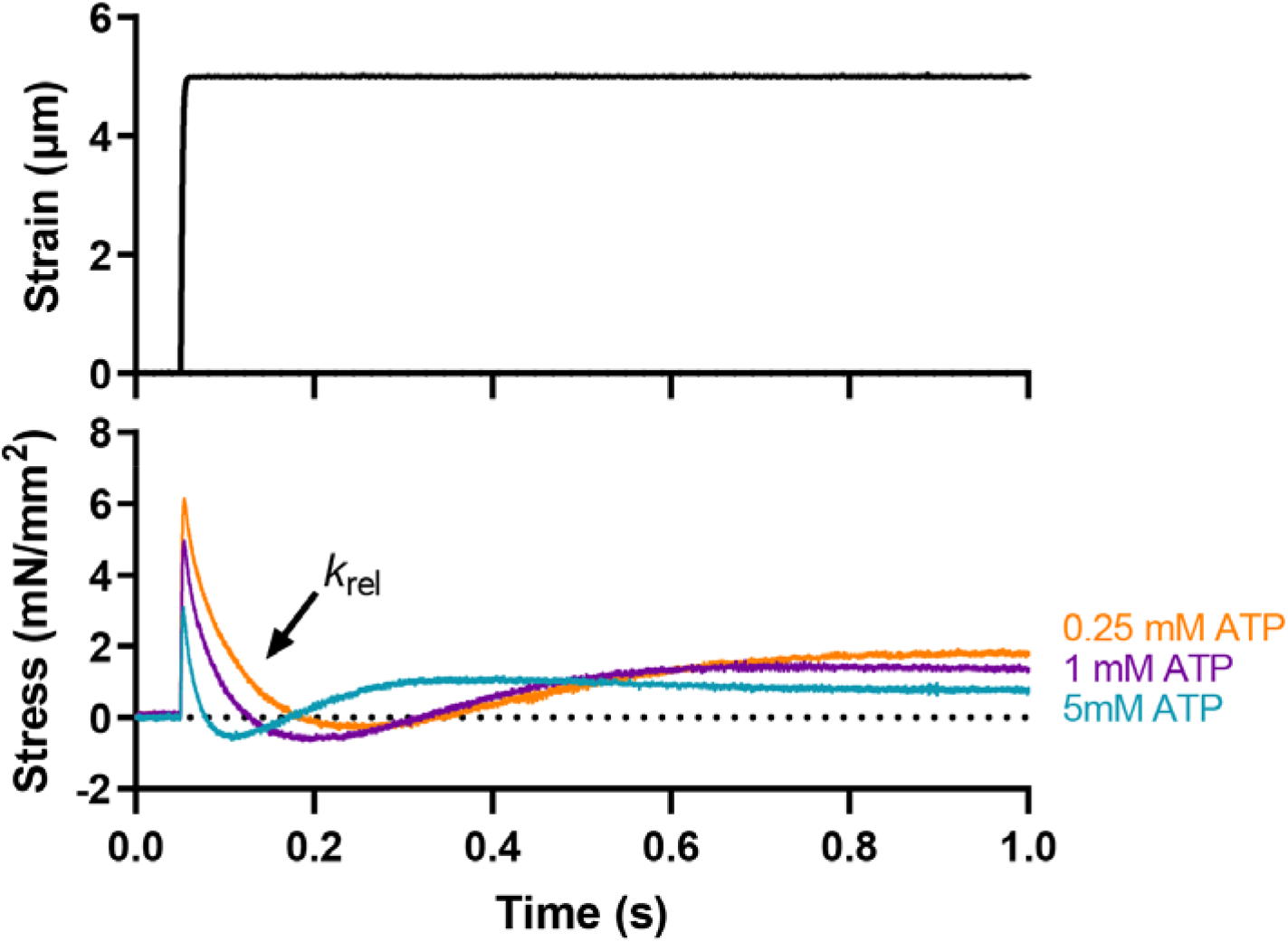
Quick stretch protocol used for calculating nucleotide binding and release rates. Stress (=force per cross-sectional area) responses were recorded following a step-length change of 0.5% muscle length (above) to assess cross bridge kinetics with increasing [MgATP]. The stress response was fit to a dual exponential function to calculate the rate of stress release (*k*_rel_) associated with the cross-bridge detachment rate.

**Figure S7.**
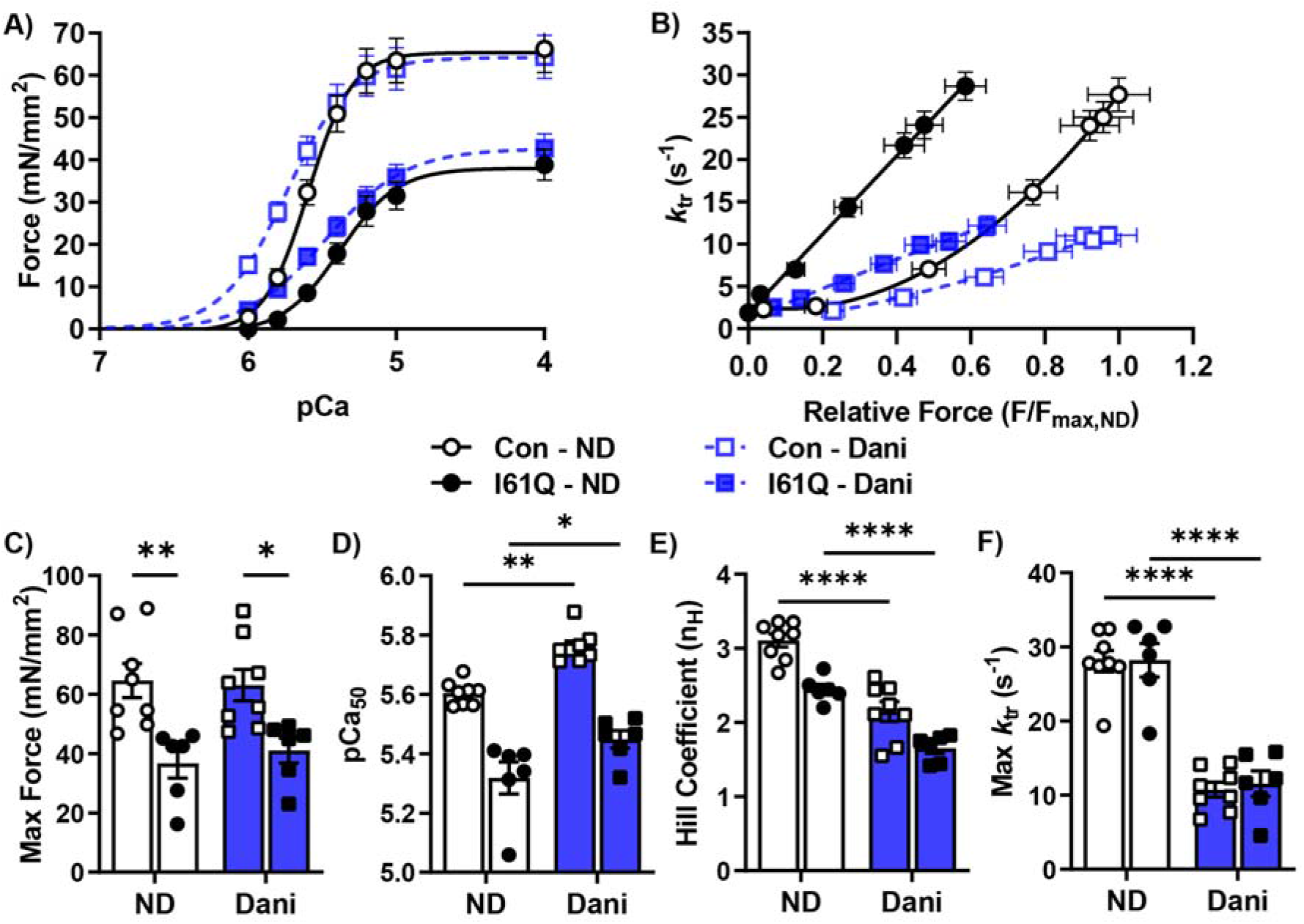
Permeabilized mechanics in control and I61Q cTnC mouse cardiac tissue treated with 1 μM Danicamtiv show increased force and calcium sensitivity. Force vs. pCa curves (A) show a right shift in I61Q cTnC compared to control mice. Treatment with 1 μm Dani results in a left shift in both control and I61Q cTnC hearts. Force redevelopment rate constants (*k*_tr_) plotted against relative force (B) was steeper in I61Q cTnC hearts and flattened in both genotypes after treatment with Dani. Maximum force was reduced in I61Q cTnC compared to control and not increased in either group after treatment with Dani (C). Treatment with Dani increased pCa_50_ in both genotypes(C) with a significant decrease in Hill coefficient (D) and max *k*_TR_ (E).

**Figure S8.**
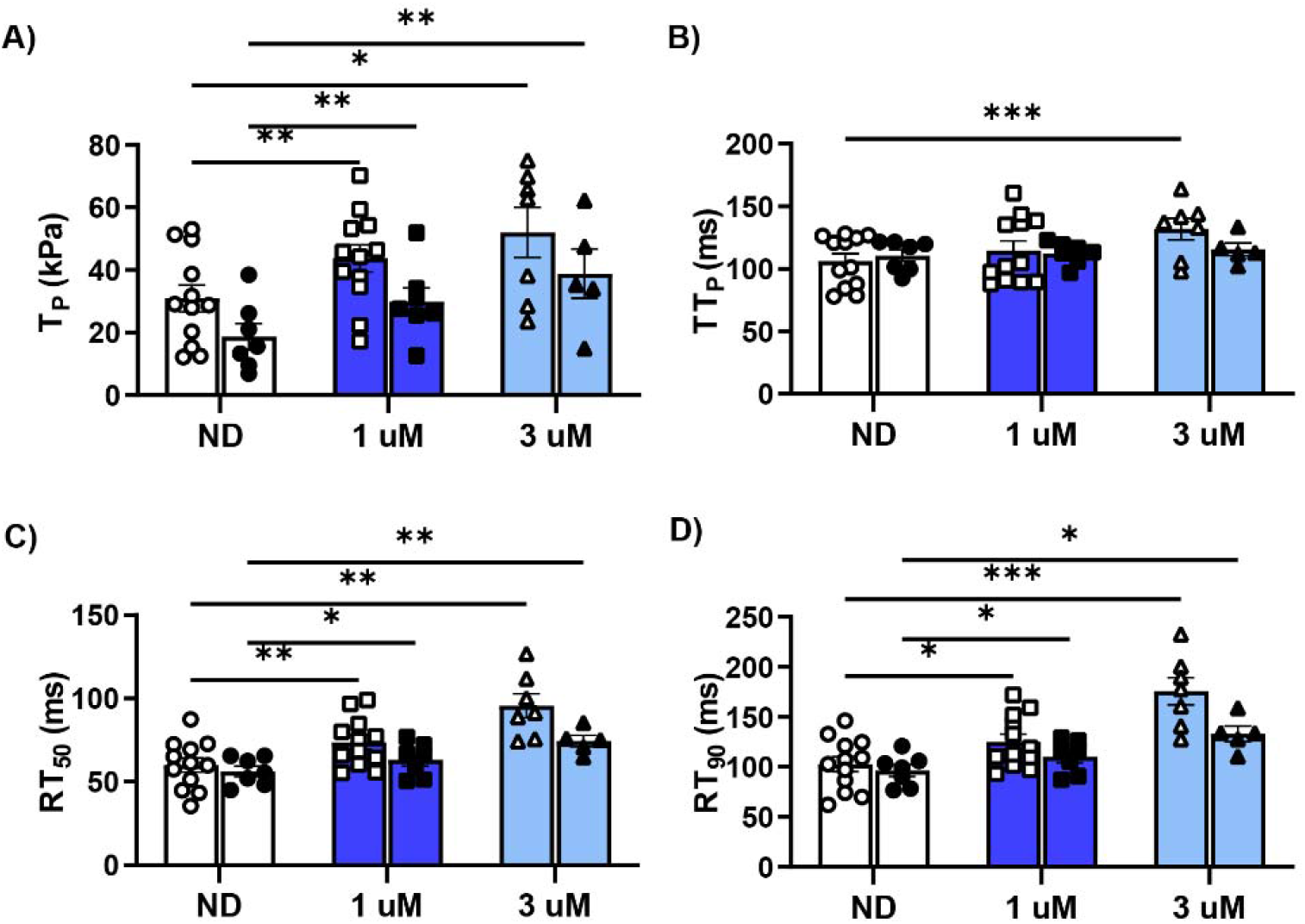
Danicamtiv increased peak tension (T_p_) and prolonged relaxation in both control and I61Q cTnC cardiac twitches. T_p_ is reduced in I61Q cTnC compared to control mice and both increased after treatment with Dani in dose dependent manner (A). Time to peak (TT_P_) was not affected at baseline with slight increase at 3 μm Dani in control mice only (B). Time to 50% (RT_50_) and 90% (RT_90_) relaxation time were similar in both genotypes at baseline. Treatment with 1 and 3 μm Dani increased both relaxation times with a larger effect in control hearts.

**Figure S9.**
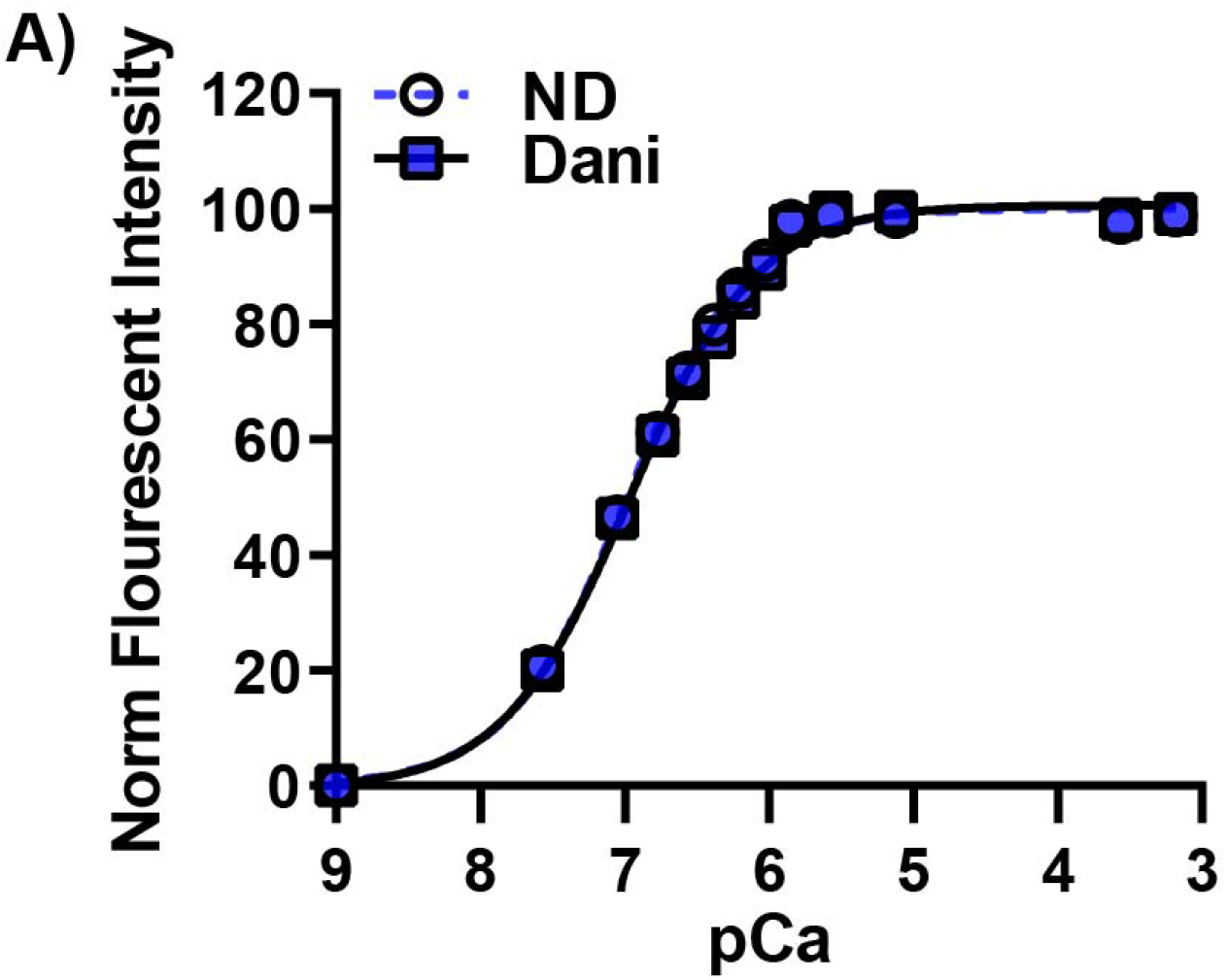
1 μM Danicamtiv did not alter calcium binding to cTnC. Dani did not alter Ca^2+^-dependent changes in the fluorescence of recombinant human 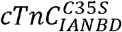. Excitation was at 490 nm, and emission was monitored at 530 nm. Data is represented as fluorescence normalized to the maximum for each experiment.

